# Aub, Vasa and Armi localization to phase separated nuage is dispensable for piRNA biogenesis and transposon silencing in *Drosophila*

**DOI:** 10.1101/2023.07.25.549160

**Authors:** Samantha Ho, Nicholas P Rice, Tianxiong Yu, Zhiping Weng, William E Theurkauf

## Abstract

From nematodes to placental mammals, key components of the germline transposon silencing piRNAs pathway localize to phase separated perinuclear granules. In *Drosophila,* the PIWI protein Aub, DEAD box protein Vasa and helicase Armi localize to nuage granules and are required for ping-pong piRNA amplification and phased piRNA processing. *Drosophila* piRNA mutants lead to genome instability and Chk2 kinase DNA damage signaling. By systematically analyzing piRNA pathway organization, small RNA production, and long RNA expression in single piRNA mutants and corresponding *chk2/mnk* double mutants, we show that Chk2 activation disrupts nuage localization of Aub and Vasa, and that the HP1 homolog Rhino, which drives piRNA precursor transcription, is required for Aub, Vasa, and Armi localization to nuage. However, these studies also show that ping-pong amplification and phased piRNA biogenesis are independent of nuage localization of Vasa, Aub and Armi. Dispersed cytoplasmic proteins thus appear to mediate these essential piRNA pathway functions.

## Introduction

24-31 nucleotide long piRNAs are bound to PIWI clade Argonaute proteins and have a conserved function in silencing transposons in the germline (Aravin, Hannon, & Brennecke, 2007; Brennecke et al., 2007; Siomi, Sato, Pezic, & Aravin, 2011; Vagin et al., 2006). In *Drosophila,* the most abundant germline piRNAs originate from pericentromeric and subtelomeric “clusters”, composed of nested transposon fragments (Bergman, Quesneville, Anxolabehere, & Ashburner, 2006; Brennecke et al., 2007). These heterochromatic loci are bound by the Rhino-Deadlock-Cutoff complex (RDC), which promotes transcription from both genomic strands and suppresses splicing and polyadenylation, producing long unspliced piRNA precursors (Chen et al., 2016; Klattenhoff et al., 2009; Mohn, Sienski, Handler, & Brennecke, 2014; Z. Zhang et al., 2014). UAP56 and the THO complex are components of the TREX, which binds to piRNA precursors and facilitates nuclear export by the noncanonical Nxf3-Nxt1 mediated Crm1 pathway (ElMaghraby et al., 2019; Hur et al., 2016; Kneuss et al., 2019; Mendel & Pillai, 2019; F. Zhang et al., 2012; G. Zhang et al., 2018). Following nuclear export, the conserved DEAD box RNA binding protein Vasa and the PIWI clade Argonautes Aub and Ago3, which colocalize to perinuclear nuage granules, drive ping-pong piRNA processing of precursor and transposon transcripts (Brennecke et al., 2007; Lim & Kai, 2007; Webster et al., 2015; Xiol et al., 2014). Ago3 catalyzed cleavage also generates precursors for phased piRNA biogenesis (Ge et al., 2019). In this branch of the pathway, processive processing of long precursors produce head-to-tail piRNAs that are loaded into Piwi, which localize to the nucleus, and guide transcriptional silencing (Cox, Chao, & Lin, 2000; Kalmykova, Klenov, & Gvozdev, 2005; Klenov et al., 2011). Phased biogenesis requires the putative helicase Armi and endonuclease Zuc, and Armi localizes to nuage and mitochondria, and Zuc localizes to the outer mitochondrial membrane (Ge et al., 2019; Han, Wang, Li, Weng, & Zamore, 2015; Mohn, Handler, & Brennecke, 2015). These findings suggest that piRNA precursors transit from clusters to nuage, where they undergo ping-pong biogenesis or are transferred to mitochondria for phased biogenesis.

Supporting this model, we previously found that prominent perinuclear nuage granules containing Aub and Vasa localize across the nuclear envelope from piRNA clusters marked by the RDC, and that Aub and Vasa are dispersed in the cytoplasm in *uap56* mutants, which disrupt piRNA precursor export from the nucleus (F. Zhang et al., 2012). Based on these findings, we proposed that export of piRNA precursors promotes assembly of the perinuclear ping-pong amplification machinery. However, *uap56* mutations lead to transposon activation, genome instability, and DNA damage signaling through Checkpoint kinase 2 (Chk2), which could disrupt nuage. To distinguish between these alternatives, we compared piRNA production, transposon silencing and nuage organization in *uap56* single mutants and *uap56* double mutants with *mnk*, which encodes the *Drosophila* homolog of Chk2. Significantly, Vasa and Aub localization to perinuclear granules is restored in *uap56, mnk* mutants. However, localization of the nuage granules that form is not biased toward regions of the nuclear envelope opposite clusters. Thus, TREX-dependent precursor export biases perinuclear nuage location, and damage signaling through Chk2 disrupts nuage association of key components of the ping-pong machinery.

Motivated by these observations, we analyzed the impact of Chk2 signaling on piRNA pathway organization, transposon silencing and gene expression in mutations that disrupt cluster transcription (*rhi*), piRNA precursor export from the nucleus (*thoc7*), ping-pong amplification (*aub, ago3, vasa*), and phased biogenesis (*armi*). These studies revealed a primary function for Rhino-dependent piRNA precursor transcription in localization of the ping-pong machinery to nuage, and co-dependencies between Aub, Ago3 and Vasa for localization to these perinuclear granules.

Vasa was the first molecularly identified nuage component and is considered essential to nuage assembly and ping-pong piRNA amplification, and localizes to the posterior pole of the developing oocyte and early embryo and is required for embryonic axis specification (Lasko & Ashburner, 1990; Liang, Diehl-Jones, & Lasko, 1994; Xiol et al., 2014). Axis specification and ping-pong amplification are not restored in *mnk, vas* double mutants, consistent with primary functions for Vasa in both processes. However, we also find that Ago3 localizes to perinuclear granules, and that electron dense nuage persists, in *vasa* null mutants. We also show that a previously characterized *vas* mutation that disrupts embryonic axis specification has minimal impact on piRNA production or transposon silencing. This separation-of-function mutation produces very low levels of wild type Vasa protein, which appears to be expressed exclusively during the earliest stages of oogenesis (Lasko & Ashburner, 1990). In this mutant background, Vasa is undetectable and Aub does not localize to nuage from stage 1 through 14 of oogenesis. Vasa thus appears to initiate ping-pong amplification, but is not required for nuage assembly or propagation of the ping-pong cycle though most of oogenesis. These findings reveal a novel function of Chk2 in control piRNA pathway organization, but also indicate the piRNA biogenesis and transposon silencing are catalyzed by proteins that are dispersed in the cytoplasm. We reconsider nuage function in light of these findings.

## Results

### *Uap56* mutations trigger Chk2-modification of perinuclear nuage

In *Drosophila melanogaster,* germline piRNA precursors are exported from the nucleus and undergo ping-pong processing by the DEAD box protein Vasa and PIWI proteins Aub and Ago3 (Brennecke et al., 2007; Xiol et al., 2014). All three ping-pong proteins localize to electron dense nuage granules, which are closely associated with the nuclear envelope (Liang et al., 1994; Lim & Kai, 2007). Uap56 and the heteropentameric THO complex are components of the <u>tr</u>anscription and <u>ex</u>port complex (TREX), which mediates nuclear export of piRNA precursors (Hur et al., 2016; Rehwinkel et al., 2004; G. Zhang et al., 2018). The *uap56^sz15^* point mutation alters a conserved surface residue and reduces UAP56 binding to THO (G. Zhang et al., 2018). This allele is homozygous lethal, but viable and female sterile in combination with *uap56^28^*, which produces low levels of wild type protein (Meignin & Davis, 2008). The *uap56^sz15/28^* combination does not alter gene expression, but disrupts transposon silencing, piRNA biogenesis, and Aub and Vasa localization to perinuclear granules (F. Zhang et al., 2012). Uap56 and THO components co-localize with piRNA clusters marked by the Rhino-Deadlock-Cuff complex (RDC), and nuage granules localize across the nuclear envelope from piRNA clusters at the nuclear periphery (F. Zhang et al., 2012; G. Zhang et al., 2018). Based on these findings, we proposed that UAP56-dependent piRNA precursor export promotes assembly of perinuclear nuage.

The *uap56^sz15/28^* combination leads to transposon overexpression, DNA damage, and activation of Checkpoint Kinase 2 (Chk2), which could indirectly disrupt nuage. To distinguish between a direct role for Uap56/TREX in nuage organization and Chk2-dependent disruption of this structure, we localized nuage components in *uap56^sz15/28^* mutants and *uap56^sz15/28^* double mutant with a null allele of *mnk,* which encodes the *Drosophila* Chk2 homolog. For this analysis, we evaluated localization of the ping-pong factors Vasa, Aub and Ago3 and the putative helicase Armi, which promotes phased piRNA biogenesis and localizes to nuage and mitochondria. To objectively quantify protein localization, we used laser scanning confocal imaging to optically section labeled egg chambers and computationally defined nuage as foci adjacent to the nuclear envelope with signal four standard deviations above average signal for the image (see Methods).

In control egg chambers, double label fluorescence imaging indicates that Aub and Ago3 colocalize with Vasa in prominent perinuclear foci (filled arrows, Figure 1A-F), and that Aub and Ago3 show essentially identical distributions (Figure S1D-H). Confirming these qualitative observations, quantitative imagine shows that nuage signal for Aub and Ago3 broadly correlate with Vasa signal (Figure 1G-J). Armi also co-localizes with Vasa in perinuclear nuage (Figure S1A-C) (Ge et al., 2019), indicating that Vasa, Aub, Ago3 and Armi are components of that same prominent nuage foci. In *uap56^sz15/28^* mutants, by contrast, direct inspection and quantification of fluorescent signal shows that Aub is dispersed in the cytoplasm (Figure 2E and 2K) and Vasa and Ago3 localization to perinuclear granules is reduced (Figure 2B and K, S2B, S2E and S2K). In striking contrast, nuage localization of Aub, Ago3 and Vasa are restored in *mnk, uap56^sz15/28^* (Figure 2C, 2F, 2L, Figure S2C, S2F, 2SL). The *uap56* mutation thus triggers Chk2-dependent loss of ping-pong biogenesis factors from perinuclear granules. Conversely, Armi does not localize to nuage in *uap56^sz15/28^* single or *mnk* double mutants (Figure S2M), indicating that UAP56 has a more direct role in Armi localization to nuage.

**Figure 1:**
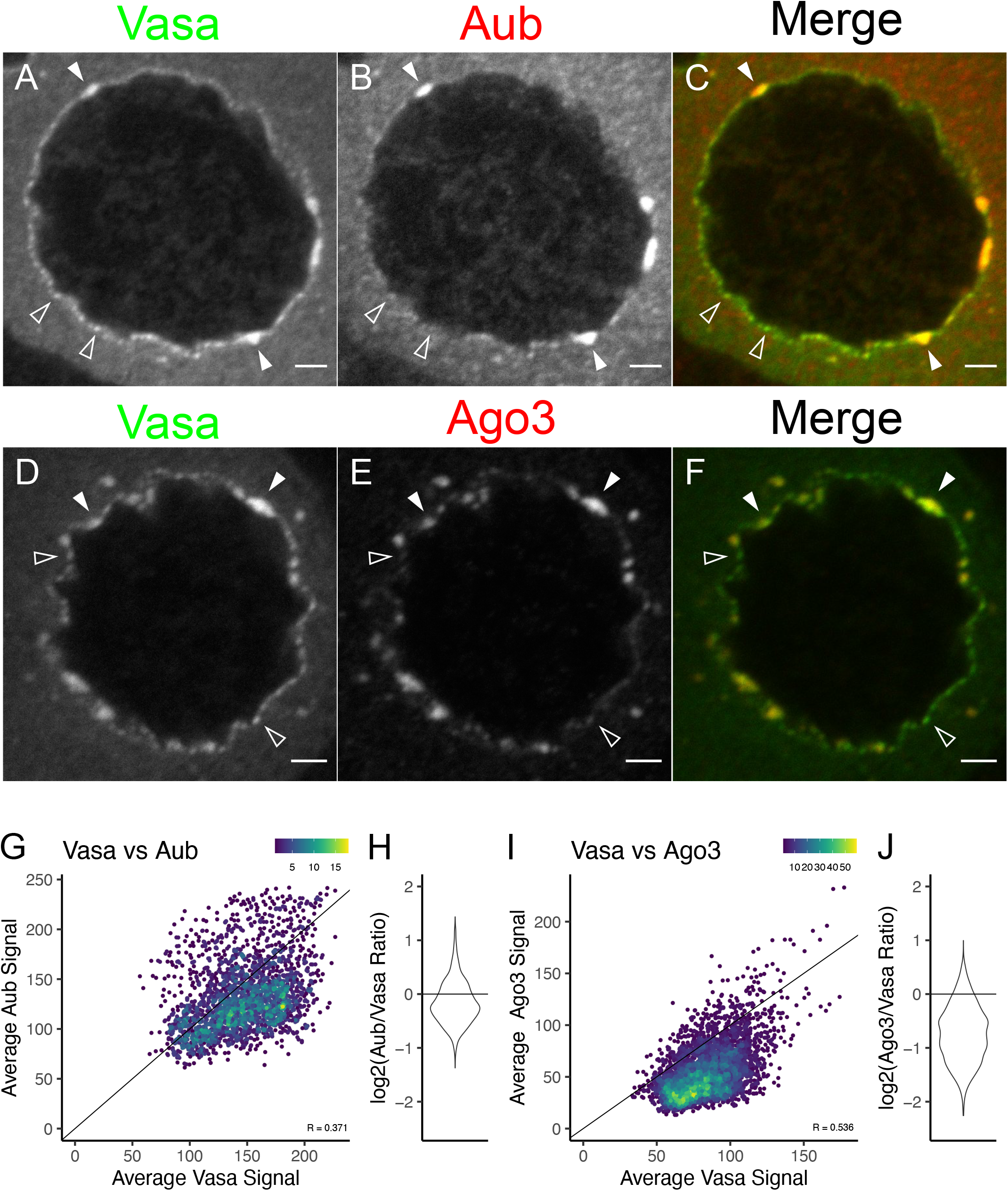
Co-localization of the ping-pong machinery to prominent nuage granules. A-C. Confocal images of *w^1^* nurse cell nucleus double labeled for Vasa (green) and Aub (red) show colocalization of the two nuage markers (filled arrow heads) along with instances where there is Vasa localize without Aub (open arrow heads). Scale bar is 2 μm. D-F. Confocal images of *w^1^* nurse cell nucleus double labeled for Vasa (green) and Ago3 (red) show similar trends as above. G. Density scatterplot comparing of average Vasa signal vs average Aub signal where each point represents a nuage granule in *w^1^*. Only granules proximity to nuclear pore signal was used for this analysis. Pearson correlation coefficient was calculated for each comparison. H. Violin plot showing distribution of Aub/Vasa signal ratios where 0 means equal amount of both signals, positive is bias towards more Aub signal and negative values are bias towards more Vasa signal. I and J. Same image quantification analysis as G and H respectively except using Ago3 instead of Aub as the second nuage marker.

**Figure 2:**
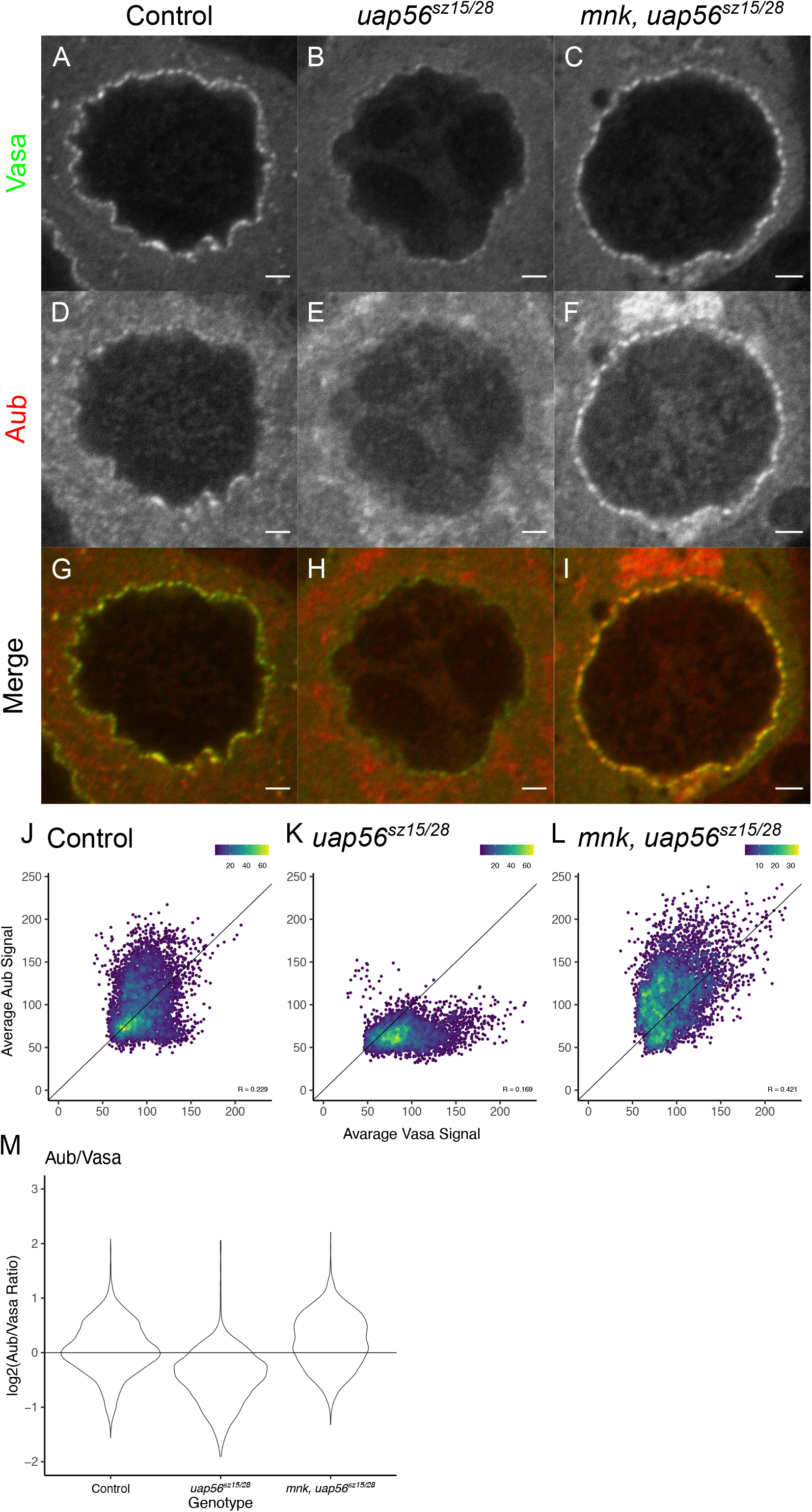
Chk2 regulates perinuclear nuage composition in *uap56* piRNA mutant. A-I. Immunofluorescence for Vasa (green) and Aub (red), in heterozygous controls, *uap56^sz15/28^*, and *mnk, uap56^sz15/28^* double mutants. Scale bar is 2 μm.

J-L. Scatterplot comparing of average Vasa signal vs average Aub signal where each point represents a perinuclear nuage granule for heterozygous controls*, uap56^sz15/28^*, and *mnk, uap56^sz15/28^* double mutants respectively.

M. Violin plot showing distribution of Aub/Vasa signal ratios.

The defects in Aub, Ago3, Vasa and Armi localization in *uap56* mutants could reflect loss of nuage, or reduced localization of these proteins to nuage granules. To distinguish between these alternatives, we used transmission electron microscopy (EM) to directly assay for nuage in *uap56^sz15/28^* and *mnk, uap56^sz15/28^* mutants. As shown in Figure 3A-3C, nuage is present in both genetic backgrounds. Chk2 activation thus displaces Vasa, Aub, and Ago3 from nuage but does not block nuage assembly.

**Figure 3:**
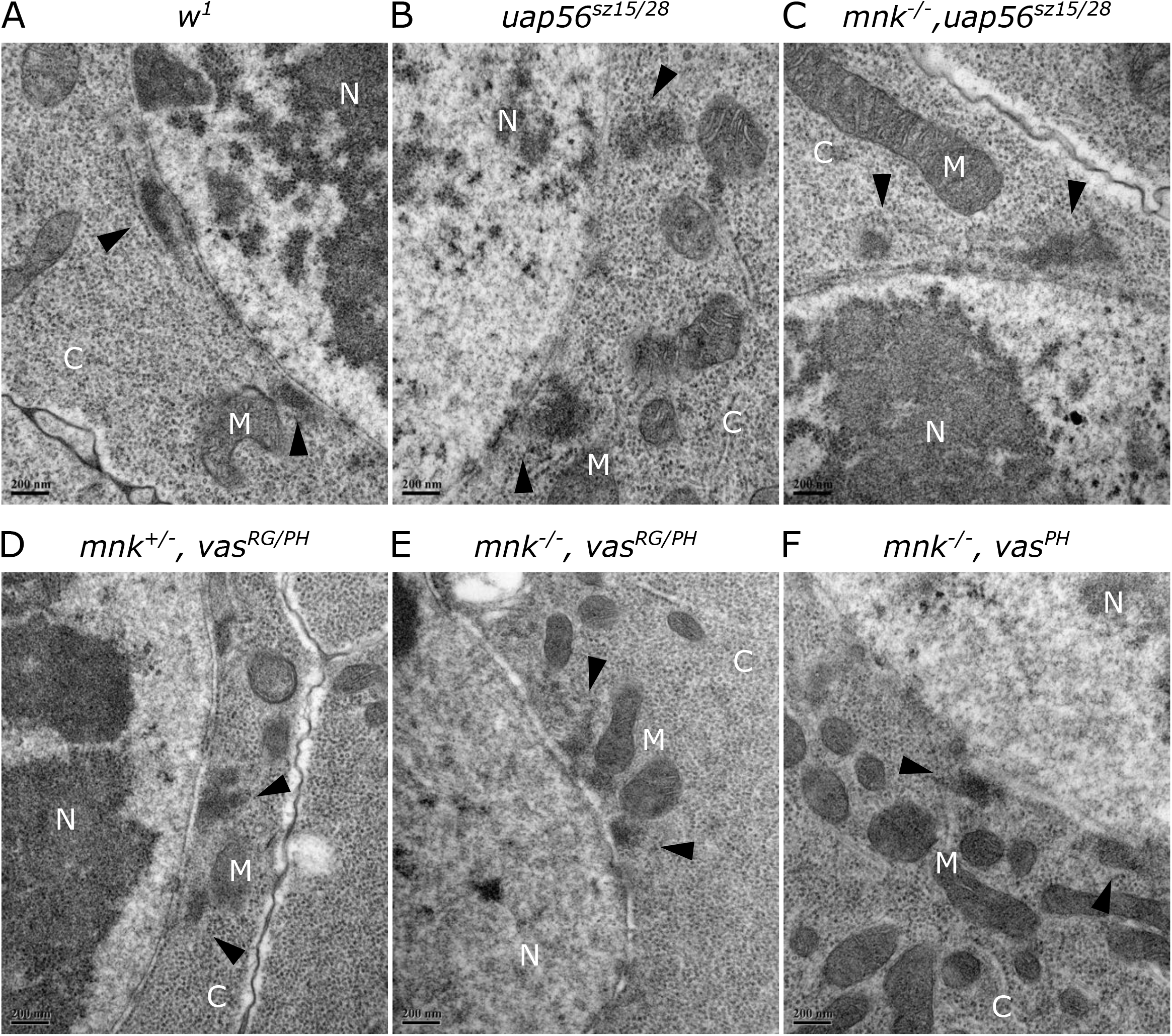
Electron dense nuage granules detected in various piRNA mutants. A. Electron microscopy (EM) images of Stage 3-5 nurse cells for WT (*w^1^*) control along with various piRNA mutants show electron dense nuage structures designated by the black arrow heads. N (nucleus), C (cytoplasm), and M (mitochondria). B and C. There are no detectable differences in nuage structure between *uap56^sz15/28^* and *mnk^-/-^, uap56^sz15/28^* double mutants. D and E. The *vas^RG/PH^* allelic combination produce tiny amounts of Vasa early on during oogenesis but is undetectable in the stages imaged still show intact nuage structure via EM. *mnk^-/-^*, *vas^RG/PH^* double mutants also intact nuage granules. F. *vas^PH^* homozygotes are a compete deletion of the open reading frame of Vasa still show intact electron dense nuage structure via EM.

### Many piRNA pathway proteins are dispensable for nuage assembly

The conserved DEAD box protein Vasa is reported to have an essential role in nuage assembly (Liang et al., 1994), but nuage persists in *uap56^sz15/28^* mutants, despite loss of Vasa from perinuclear granules. Low levels of Vasa at nuage could be sufficient to promote granule assembly, or Vasa could be dispensable for nuage formation. We therefore used transmission electron microscopy to analyze ovaries isolated from a *vas* null allelic combination, *vas^RG/PH^* and in the corresponding *mnk* double mutant. In addition, we analyzed ovaries homozygous for a *vas* deletion (*vas^PH^*) by looking at the *mnk, vas^PH/PH^* double mutants (Figure 3D-F). Electron dense perinuclear granules are present in all of these backgrounds. In addition, immunofluorescence labeling shows that Ago3 and Armi localize to perinuclear granules in *vas^RG/PH^* null ovaries (Figure S3C and S3D). Nuage assembly is therefore independent of Vasa. However, Aub does not localize to nuage in *vas^RG/PH^* mutants or *mnk, vas^RG/PH^* double mutants (Figure S3B), indicating that Vasa is required to recruit this PIWI protein to nuage.

To determine if other piRNA pathway genes are required for nuage assembly, we used thin section transmission electron microscopy to analyze *rhi^KG/02^*, *thoc7^d/Df^*, *aub^HN/QC^*, and *mnk; armi^1/72.1^* mutant ovaries (Figure S4). As observed in *uap56* and *vas* mutants, electron dense perinuclear nuage is present in all of these backgrounds. In addition, immunofluorescence imaging show that Ago3 also localizes to perinuclear granules in almost all of these mutant backgrounds (Figure S3C). Nuage assembly thus does not require Rhi, Vasa, Aub, Ago3 or Armi.

### piRNA production and transposon silencing

Localization of piRNA pathway proteins to perinuclear granules is deeply conserved, implying an important role in biogenesis and transposon silencing (Aravin et al., 2009; Batista et al., 2008; Lim & Kai, 2007). The *uap56^sz15/28^* allelic combination reduces piRNA expression and ping-pong amplification (Figure 4A and B, (F. Zhang et al., 2012)). To determine if these defects are the result of Chk2-dependent loss of Aub and Vasa localization to nuage, we sequenced small RNAs from *uap56^sz15/28^* and *mnk, uap56^sz15/28^* double mutant ovaries, and *w^1^* and *mnk* controls. Since *w^1^* and *mnk* controls show similar levels of piRNAs mapping to transposon, ping-pong z-scores, transposon expression, and cluster expression (Figure S5), all subsequent comparisons use *w^1^* as WT control, unless otherwise stated. As shown in the scatter plots in Figure 4A, anti-sense transposon mapping piRNA levels in *uap56^sz15/28^* and *mnk, uap56^sz15/28^* double mutants are significantly reduced relative to *w^1^* controls. However, direct comparison of *uap56^sz15/28^* and *mnk, uap56^sz15/28^* indicate that anti-sense piRNA levels are approximately two-fold higher than in the double mutants. The z-score measuring enrichment of the 10 nucleotide overlap between sense and antisense transposon mapping piRNAs also show a slight increase in the double mutants (Figure 4B). Uap56 thus has a primary function in piRNA biogenesis, while Chk2-dependent loss of the central ping-pong amplification factors Aub and Vasa has a remarkably modest impact on the piRNA biogenesis in the *uap56* mutant background.

**Figure 4:**
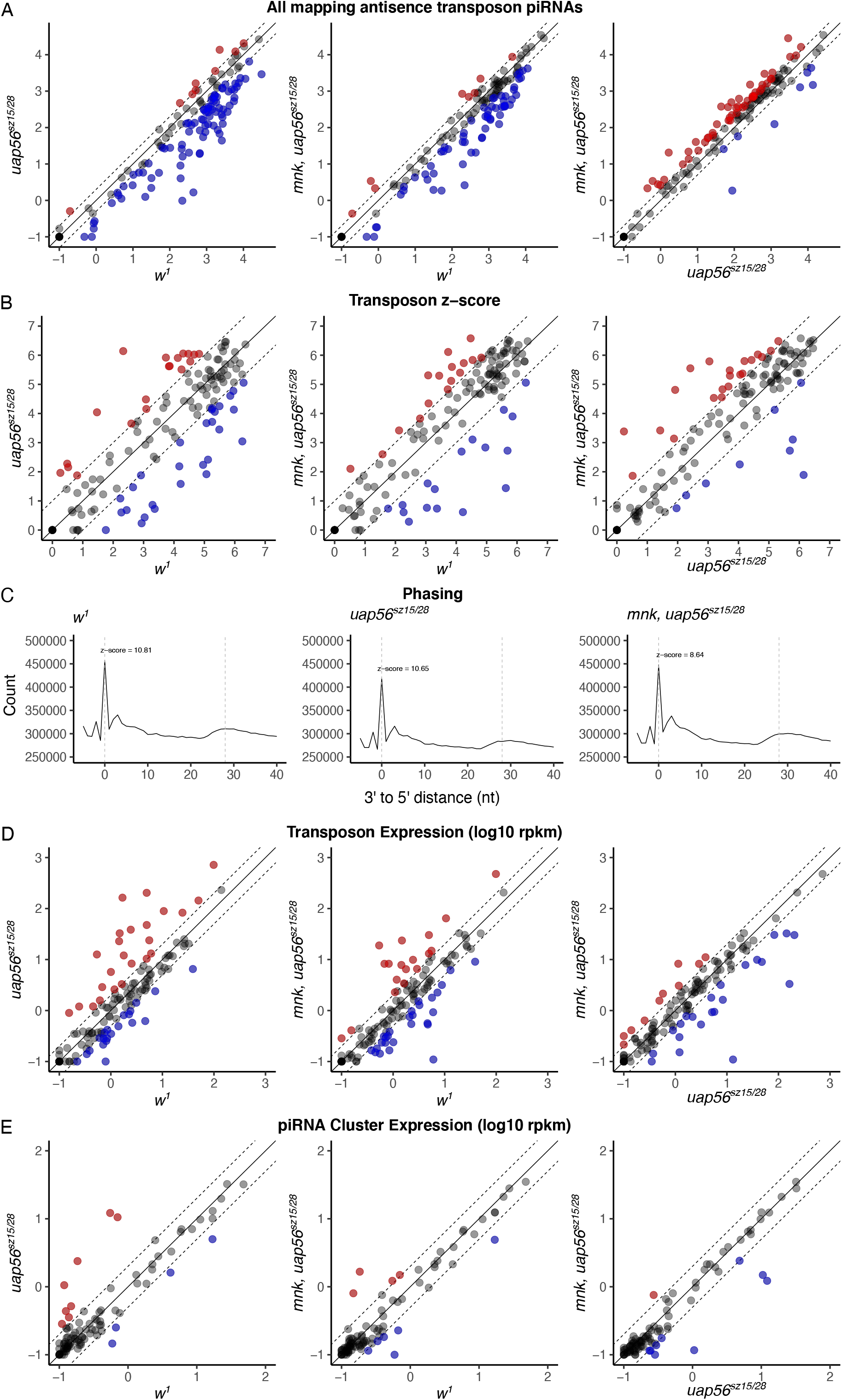
Chk2 activation show subtle effects on piRNA production and transposon silencing. A. Scatterplot comparing antisense transposon mapping piRNA abundance (log 10) between *w^1^*, *uap56^sz15/28^*, and *mnk, uap56^sz16/28^* double mutants. Each point represents a transposon family. Red and blue indicate the 2-fold increase and decrease respectively. B. Scatterplot comparing pong-pong z-scores (log 2) detected for each transposon family with the same setup as above. C. Phasing was measured by looking at the distribution of the shortest distance between 3ʹ end of one piRNA and the 5ʹ end of the subsequent piRNA downstream. Zero nucleotide distance represents the head of a piRNA abutted directly downstream of the tail of another piRNA with no nucleoid separating the two. Enrichment is detected at zero nucleotide distance for *w^1^*, *uap56^sz15/28^*, and *mnk, uap56^sz16/28^* double mutants which is a signature that phasing pathway is intact and functional. D. Scatter plots show pairwise comparisons of transposon expression levels (log 10 rpkm) between *w^1^* wild type control*, uap56^sz15/28^* and *mnk, uap56^sz15/28^* double mutants. Each point represents a transposon family. Red and blue indicate the 2-fold increase and decrease respectively. Transposons are over expressed in *uap56* and *mnk* doubles, however upregulation of a subset of these transposons is due to activation of Chk2. E. Scatter plots with same pairwise comparisons as above except showing piRNA cluster expression levels instead of transposon expression. piRNA cluster expression is less sensitive to activation of Chk2.

The Armi helicase promotes processive piRNA production during phased biogenesis and localizes to nuage and mitochondria (Ge et al., 2019). Ago3 localizes to nuage, and products of Ago3 cleavage undergo phased processing by the mitochondrial nuclease Zuc (Han et al., 2015; Mohn et al., 2015; Wang et al., 2015). Together, these findings suggest that Armi shuttles precursor transcripts from nuage to mitochondria. Armi does not localize to nuage in *uap56^sz15/28^* or *mnk, uap56^sz15/28^*. To analyze phasing in these mutants, we quantified the shortest nucleotide distance between the 3ʹ ends of one piRNA and the 5ʹ end of the downstream piRNA. A primary peak at 0 nucleotides and secondary peak at 28 nucleotides, which are hallmarks of phased processing, are observed for w^1^ controls, *uap56^sz15/28^* and *mnk, uap56^sz15/28^* (Figure 4C). Phased piRNA biogenesis thus appears to be independent of UAP56, and does not require Armi accumulation at nuage. However, Armi could transiently localize to perinuclear granules in *uap56* mutants. In addition, the *uap56^sz15/28^* combination is hypomorphic and produces low levels of wild type proteins, which could be sufficient to promote phased piRNA biogenesis.

To determine the impact of Chk2 on transposon silencing and piRNA precursor expression, we performed long RNA sequencing on *uap56^sz15/28^* and *mnk, uap56^sz15/28^* double mutants. A few piRNA clusters show increased expression and others show decreased expression in *uap56^sz15/28^* and *mnk, uap56^sz15/28^* relative to *w^1^,* but the majority of clusters are expressed at comparable levels in all of these backgrounds (Figure 4E). A subset of transposons families are significantly over-expressed in both *uap56^sz15/28^* and *mnk, uap56^sz15/28^* double mutants relative to *w^1^,* supporting a direct function for UAP56 in the piRNA pathway (Figure 4D, red points). By contrast, transposon expression is comparable in *mnk, uap56^sz15/28^* double mutants and *uap56^sz15/28^* single mutants (Figure 4D). Within the genetic context of *uap56* mutants, nuage localization of Aub and Vasa thus have a surprisingly modest impact on piRNA biogenesis and transposon silencing.

### Uap56 is required for spatial juxtaposition of nuage and clusters

Nuage granules are biased toward regions of the nuclear envelope opposite piRNA clusters, and *uap56* mutations disrupt TREX-mediated cluster transcript export from the nucleus and disrupt Aub and Vasa localization to nuage, leading us to propose that piRNA precursor export organizes perinuclear nuage. However, as shown here, loss of Aub and Vasa localization in *uap56* mutants is an indirect consequence of Chk2 signaling. We therefore quantified the spatial relationship between clusters and perinuclear granules in control, *uap56^sz15/28^*, and *mnk, uap56^sz15/28^* double mutants. For this analysis, we triple labeled egg chambers for clusters with anti-Rhino, the nuclear envelope with a pan-nuclear pore antibody, and for nuage with antibodies to Aub or Ago3. Triple labeling for Rhino, nuclear pore and Armi was also performed as a negative control, since Armi does not localize in the *uap56^sz15/28^* or *mnk, uap56^sz15/28^*. Labeled egg chambers were optically sectioned by laser scanning confocal microscopy (Figure 5A-I), and clusters and nuage were defined as foci with signal three standard deviations above average signal. Clusters at the nuclear periphery were identified by proximity to the nuclear envelope, and the fraction of clusters at the periphery with nuage directly across the nuclear envelope, termed percent juxtaposed, was calculated (see Methods).

**Figure 5:**
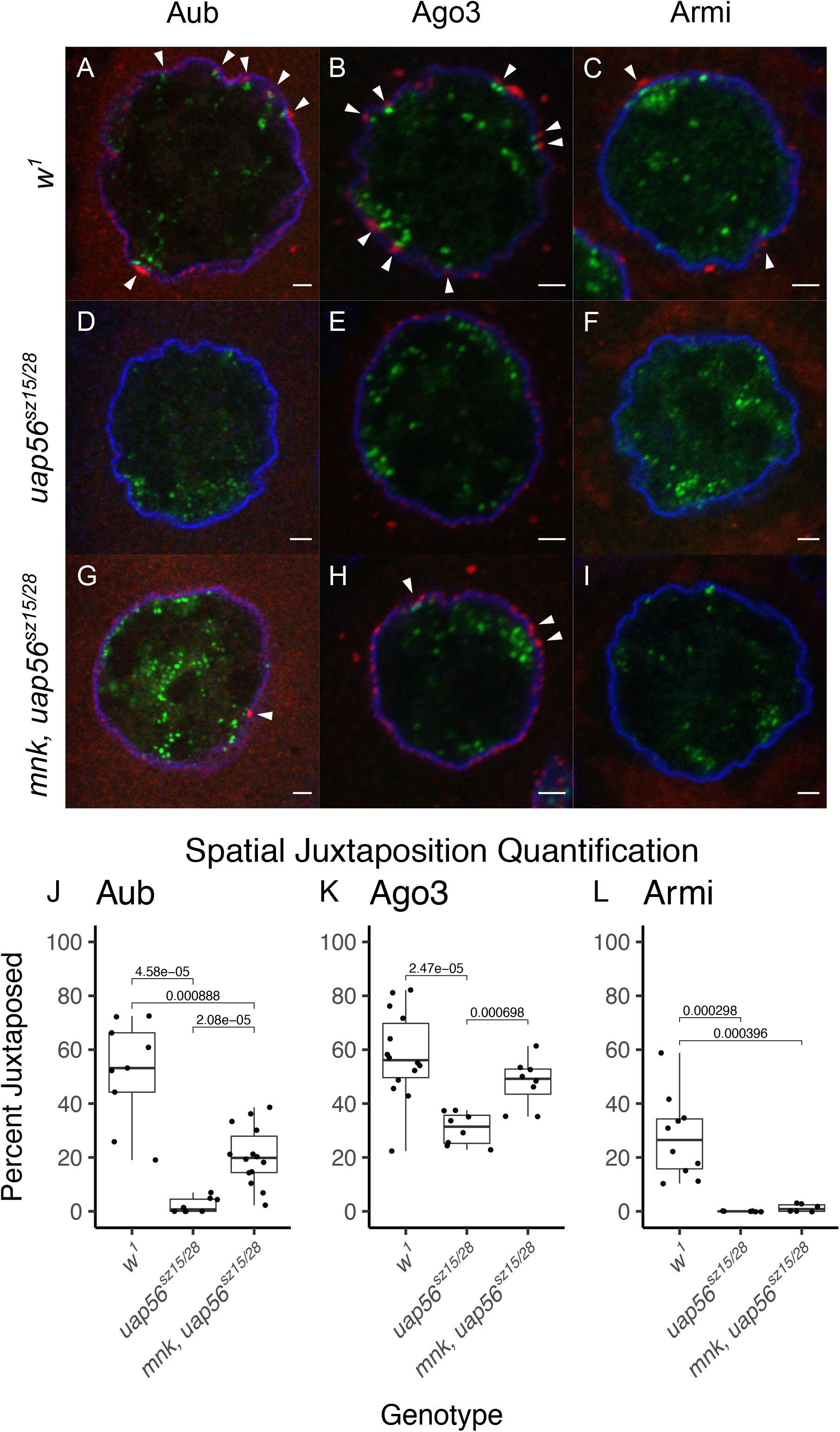
UAP56 is required for Armi nuage localization and organizing nuage adjacent to nuclear piRNA machinery. A-C. Confocal immunofluorescence images of WT control (w1) nurse cell nucleus stage 6-8 triple labeled for Rhino (green), nuclear pore (blue) and a nuage marker (red), Aub, Ago3, or Armi. White arrowheads show instances where peripheral Rhino is juxtaposed to cytoplasmic nuage marker. D-F. *uap56^sz15/28^* stained the same as WT show no Aub and Armi localization and little to no detectable juxtaposition events between nuage marker and Rhino. G-I. *mnk, uap56^sz15/28^* stained the same as WT show partial restoration of juxtaposing for Aub and Ago3, however localization of Armi remains perturbed. J-L. Images from the staining experiments above were quantify for percent of peripheral rhino juxtaposed to Aub, Ago3 or Armi (see methods). Box plot show the percent juxtaposed distribution for *w^1^*, *uap56^sz15/28^*, and *mnk, uap56^sz15/28^*, where each point represents a nucleus. P-value shown whenever significant different.

In *w^1^* control egg chambers, 50 to 60% of clusters at the nuclear periphery are juxtaposed to nuage marked by Aub or Ago3 (Figure 5A, 5B, 5J and 5K). In *uap56^sz15/28^* single mutants, Aub and Ago3 nuage localization is lost and juxtaposition collapses (Figure 5D, 5E, 5J, and 5K). In *mnk, uap56^sz15/28^* double mutants, Aub localization to nuage is restored, but the fraction of clusters adjacent to nuage is significantly below control level (Figure 5G and 5J). Ago3 localization to perinuclear foci is restored in *mnk, uap56^sz15/28^* and the fraction of foci juxtaposed to clusters is reduced, but not significantly different than controls (Figure 5K). However, perinuclear Ago3 foci are numerous and uniformly distributed around the nucleus (Figure 5H), and tight juxtaposition is not restored (compare Figure 5B and H). The increase in “juxtaposition” is therefore likely the result of an increased probability of special overlap between Ago3 and Rhino foci. Armi does not localize in either *uap56^sz15/28^* nor *mnk, uap56^sz15/28^* double mutants and juxtaposition with Rhino is undetected in both genotypes (Figure 5F, 5I and 5L). These observations indicate that UAP56, and presumably precursor export, biases localization of the perinuclear ping-pong processing machinery to regions of the nuclear envelope opposite piRNA clusters.

### Chk2 modification of piRNA pathway mutations

Our analysis of *uap56* mutants demonstrated that Chk2 signaling can compromise piRNA pathway organization and has the potential to mask primary phenotypes associated with any mutation that destabilizes the germline genome, including essentially all piRNA pathway mutations. To better define the primary functions of piRNA pathway genes, we therefore extended our studies to a series of single and *mnk* double mutations in genes that encode nuclear (*rhi, thoc7*), perinuclear (*aub, ago3, vas*), and mitochondrial (*armi*) components of the piRNA biogenesis machinery. To allow for direct comparisons with published phenotypes, we analyzed well characterized allelic combinations of these genes (see methods). For each single and double mutant combination, we analyzed Aub, Ago3, Vasa and Armi localization, and sequenced small and long RNAs. A summary of phenotypes can be found in Table 1.

**Table 1:** Summary of piRNA single mutant and *mnk* double mutant phenotypes. Perinuclear nuage localization of Vasa, Aub, ago3 and Armi proteins were evaluated in various piRNA mutants and designated as more localized than wild type (+), wildtype like localization (WT), less localized than wildtype (-), not localized at all (O) and not applicable (NA). Long and small RNA sequencing were performed on various piRNA mutants and changes in global levels relative to *w^1^* controls were evaluated. Up relative to controls is designated by +, down relative to controls is designated by -, and comparable levels is designated by WT.

|  | Genotype | Nuage Localization of: |  |  |  | piRNA |  |  | Long RNA |
| --- | --- | --- | --- | --- | --- | --- | --- | --- | --- |
|  |  | Vasa | Aub | Ago3 | Armi | antisense TE | z-score | Phasing | TE |
| RDC | *rhi* KG/02 | - | O | - | - | -- | -- | WT | ++ |
|  | *mnk, rhi* KG/02 | - | O | - | - | -- | -- | WT | ++ |
| TREX<br>Export<br>Machinery | *uap56 sz16/28* | - | - | - | O | - | - | WT | ++ |
|  | *mnk, uap56 sz15/28* | WT | WT | WT | O | - | - | WT | + |
|  | *thoc7 d/Df* | - | O | WT | - | - | WT | WT | + |
|  | *mnk; thoc7 d/Df* | WT | O |  | - | - | WT | WT | + |
| Ping-Pong<br>Machinery | *vas* RG/PH | NA | O | WT | WT | - | - | WT | ++ |
|  | *mnk, vas* RG/PH | NA | O | WT | WT | - | - | WT | ++ |
|  | *aub* HN/QC | WT | NA | WT | WT | -- | -- | WT | + |
|  | *mnk, aub* HN/QC | WT | NA | WT | WT | -- | -- | WT | + |
|  | *ago3 t2/t3* | WT | O | NA | - | -- | -- | WT | ++ |
|  | *mnk; ago3 t2/t3* | WT | - | NA | - | -- | -- | WT | ++ |
| Transcriptional<br>Silencing | *armi 1/72.1* | WT |  | WT | NA | -- | -- | WT | ++ |
|  | *mnk; armi 1/72.1* | WT |  | WT | NA | -- | -- | WT | ++ |
|  | *mnk, piwi 02/ΔNLS* | ovaries were unrecoverable |  |  |  |  |  |  |  |

### Control of peri-nuclear nuage organization

RDC-mediated transcription produces piRNA precursors, the TREX mediates precursor nuclear export, and Aub, Vasa and Ago3 localize to perinuclear nuage and drive ping-pong processing of these precursors, and Armi localizes to nuage and mitochondria and functions in phased piRNA biogenesis. Mutations in *rhi* displace Vasa, Aub, Ago3 and Armi from nuage and produce a profound reduction in germline cluster transcription and piRNA production (Klattenhoff et al., 2009; Mohn et al., 2014). Significantly, these defects are not suppressed by *mnk* (Figure S3), suggesting that RDC-dependent cluster transcripts have a direct role in recruiting Vasa, Aub, Ago3 and Armi to nuage. The TREX is proposed to function downstream of Rhino in precursor transcript export (ElMaghraby et al., 2019; Hur et al., 2016; Kneuss et al., 2019; Mendel & Pillai, 2019; G. Zhang et al., 2018). However, *uap56^sz15/28^* and *thoc7^d/Df^* mutants disrupt the TREX, but show significantly less severe defects in germline piRNA production than *rhi* mutants (Figure 4A and S6A). In addition, these single mutations displace Vasa from nuage, but Vasa localization is restored in *mnk; uap56^sz15/28^* and *mnk; thoc7^d/Df^* double mutants (Figure 2A-2C and S3A). Localization of Ago3 is also independent of Uap56 and Thoc7. By contrast, Armi localization to nuage is disrupted in *uap56* and *thoc7* mutants and the corresponding *mnk* double mutants (Figure S2F, S2M and S3C-S3D). RDC-dependent transcripts thus promote Ago3, Aub, Vasa and Armi localization to nuage, but these transcripts appear to be exported by TREX-dependent and TREX independent mechanisms. The TREX-dependent pathway is required for Armi localization to nuage and “links” perinuclear nuage to piRNA clusters, but is dispensable for nuage localization of Ago3, Aub and Vasa.

Extending this analysis to mutations that disrupt ping-pong amplification revealed a series of interdependencies that determine nuage composition. Mutations in *ago3* and *vasa* displace Aub from nuage, and loss of Aub localization is not suppressed in *mnk* double mutants (Figure S3B). By contrast, Ago3 and Vasa localization are not disrupted in *aub^HN/QC^* or *mnk, aub^HN/QC^* mutants (Figure S3A and S3C). Aub localization to nuage thus required Ago3 and Vasa. However, Vasa localization is not disrupted in *ago3^t2/t3^* or *mnk; ago3^t2/t3^* mutations (Figure S3A), and Ago3 localization is not disrupted in *vasa^RG/PH^* or *mnk, vas^RG/PH^* mutants (Figure S3C). Ago3 and Vasa thus localize to nuage though independent mechanisms.

### piRNA biogenesis and transposon silencing

Key components of the ping-pong amplification machinery localize to perinuclear nuage, and we find that Chk2 signaling controls nuage localization of the ping-pong biogenesis machinery. To determine the if Chk2-dependent disruption of nuage contributes to the piRNA biogenesis and transposon silencing defects associated with the mutations analyzed here, we assayed small and long RNA expression in single mutants and the corresponding *mnk* double mutants. For each mutant pair, we quantified anti-sense transposon mapping piRNAs, ping-pong z-score, transposon and piRNA cluster expression (Figure S6 and S7).

All of the *mnk* double mutations analyzed show significant defects in piRNA expression and transposon silencing relative to control strains, and most of the mutants and *mnk* double mutants show comparable levels of anti-sense transposon mapping piRNAs, ping-pong z-score, and phased piRNA production (Figures S6). All of the genes analyzed thus have primary functions in the piRNA pathway. However, mutations in *thoc7* (Figure S6A and S6B) and *uap56* (Figure 4A and 4B), show a modest global increase in anti-sense piRNA expression and ping-pong z-score in *mnk* double mutants relative to single mutants, and this correlates with Vasa localization to nuage in the double mutants (Figure S3A). However, transposon expression is comparable in the single *and* mnk double mutants (Figure S7). Vasa and Aub localization to nuage thus appears to lead to a modest increase in piRNA biogenesis, but has little impact on transposon silencing.

Armi promotes phased piRNA biogenesis, co-localizes with Ago3 in nuage granules, and localizes to mitochondria with the endonuclease Zuc, which catalyzes phased piRNA production. Additionally, Ago3 cleavage generates precursors for phased processing (Ge et al., 2019). Together, these findings suggest that Armi shuttles precursors from nuage to mitochondria. Mutations in *rhi*, *uap56*, and *tho7* disrupt Armi localization to nuage, and nuage localization is not restored in *mnk* double mutants (Figure S2M and S3D). Significantly, phased piRNA biogenesis is comparable to wild type in *rhi, uap56* and *thoc7* single and *mnk* double mutants (Figure 4C and S6C). Armi localization to nuage thus appears to be dispensable for phased piRNA production.

### A separation of function allele of *vas*

Our findings suggest that nuage localization of Aub and Vasa has a relatively modest impact on the efficiency of ping-pong biogenesis and does not contribute to transposon silencing. However, all of the mutations analyzed compromise piRNA biogenesis, complicating interpretation of the data. By contrast, the *vas^PD^* mutation appears to uncouple ping pong amplification and transposon silencing from nuage localization of Vasa and Aub. Vasa is required for piRNA biogenesis and assembly of pole plasm, which is required for posterior patterning and germline specification during embryogenesis. The *vas^PD^* mutation over a deletion allele (*vas^PH^*) disrupts pole plasm assembly and posterior patterning (Schupbach & Wieschaus, 1986, 1991). However, we find that antisense transposon mapping piRNA expression, ping-pong amplification, transposon and piRNA cluster expression are comparable to controls (Figure 6C-E). Significantly, in *vas^PH/PD^* ovaries, Vasa protein is only detectable very early in oogenesis, prior to oocyte differentiation (Lasko & Ashburner, 1990), and Aub does not localize to nuage at any point during oogenesis (Figure 6A and B). RNAseq indicates that this allele produces very low levels of wild type *vas* transcripts (Figure 6E), consistent with expression only in the germarium. While compelling biochemical studies indicate that Vasa drives the pong-pong cycle by releasing cleaved transcripts from Piwi-piRNA complexes (Xiol et al., 2014), these observations imply that Vasa initiates the ping pong amplification during early oogenesis, but is dispensable for propagation of the cycle through most of oogenesis, and that Aub localization to nuage is not required for ping-pong amplification or transposon silencing.

**Figure 6:**
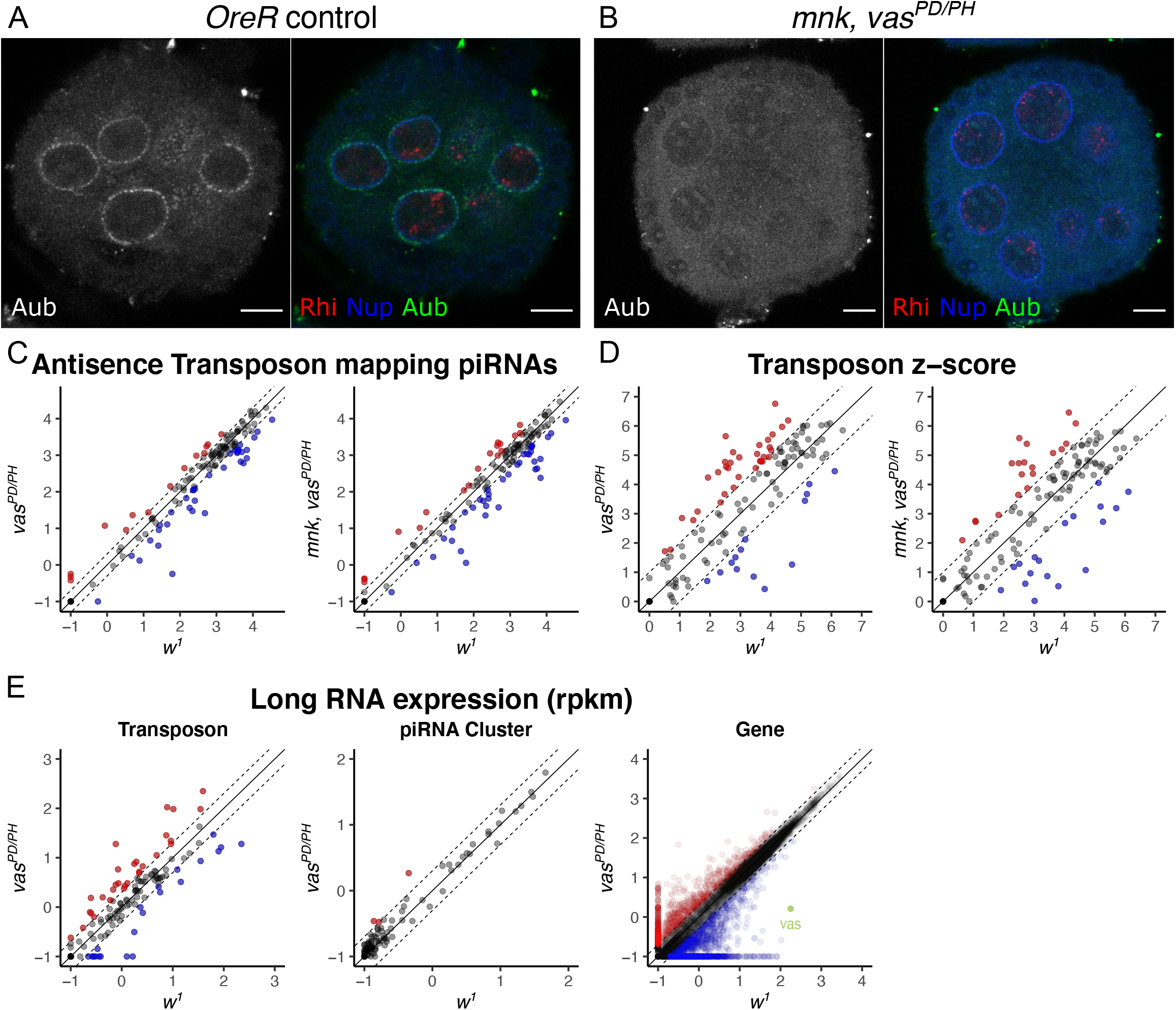
*vas^PD/PH^* allele show uncoupling of Aub localization to nuage from piRNA biogenesis, ping-pong, and transposon silencing

A-B. *OreR* control and *mnk, vas^PD/PH^* fly ovaries were stained for Aub, Nuclear pore, and Rhino. Rhino localization remains intact in *mnk, vas^PD/PH^* however Aub localization to nuage is disrupted.

C. Scatterplot show antisense transposon mapping piRNA abundance (log 10) of *vas^PD/PH^* or *mnk, vas^PD/PH^* double mutants compared to *w^1^* control. Each point represents a transposon family. Red and blue indicate the 2-fold increase and decrease respectively. piRNA production is comparable between *vas^PD/PH^* and WT controls and unaffected by activation of Chk2.

D. Scatterplot showing pong-pong z-scores (log 2) detected for each transposon family with the same comparisons as above. Transposon ping-pong z score are scattered around the diagonal suggesting ping-pong is comparable between *vasa^PD/PH^* mutants and *w^1^* controls.

E. Long RNA sequencing was performed on *vas^PD/PH^* and compared to *w^1^* controls. *vas^PD/PH^* show no global changes in transposon, piRNA cluster, or gene expression.

## Discussion

Forward genetic screens have been instrumental in defining the *Drosophila* piRNA biogenesis machinery and identifying the targets for these small silencing RNAs. However, piRNA mutations activate transposons, leading to genome instability and DNA damage signaling, complicating identification of primary phenotypes. By genetically eliminating Chk2, the genetic, cytological, and molecular studies reported here provide new insight into the primary functions of piRNA pathway genes, the role of subcellular localization in piRNA biogenesis, and the impact of DNA damage signaling on piRNA mutant phenotypes.

### DNA damage control of nuage organization

Molla-Heman et al. (2015) reported that an unusual allele of *rpp30* (*rpp30^18.1^*), which encodes the protein subunit of the tRNA processing RNaseP, blocks piRNA biogenesis and arrests oogenesis, but does not disrupt tRNA processing. Remarkably, the authors reported that the *mnk* null mutation completely suppressed the defects in piRNA expression and oogenesis, restoring fertility. These striking findings implied that DNA damage signaling through Chk2 disrupts germline transposon silencing and piRNA biogenesis. This *rpp30* allele appeared to provide a powerful tool for dissection of DNA damage control of the piRNA machinery, and the authors generously provided the relevant strains. However, following outcrossing to *w^1^* or *mnk*, we found that the *rpp30^18.1^* allele is homozygous lethal and *mnk, rpp30^18.8/PE^* females are sterile due to arrest early in oogenesis. The developmental defects associated with the *rpp30^18.1^* allele thus do not appear to result from Chk2 activation. While the allele did not prove to be useful for our studies, the published data prompted us to examine the impact of Chk2 signaling on piRNA mutants reported here. We show that Chk2 DNA damage signaling displaces various ping-pong piRNA biogenesis machinery from nuage granules.

### Nuclear control of perinuclear nuage composition

Germline piRNA biogenesis is initiated by RDC-mediated cluster transcription (Klattenhoff et al., 2009; Mohn et al., 2014). *rhi* and *mnk, rhi* mutants display a significant collapse of antisense transposon piRNAs which originate from piRNA cluster transcripts (Figure S6A). Additionally, Aub, Vasa, Ago3 and Armi are also displaced from nuage in both mutant backgrounds, consistent with a direct role for cluster transcripts in control of nuage organization (Figure S3). However, the TREX complex which mediates cluster transcript export from the nucleus, exhibits less severe defects in piRNA production than *rhi* mutants (compare Figure 4A and S6A). Additionally, both UAP56 and Thoc7 are dispensable for Vasa and Ago3 nuage localization (Figure S2 and S3). Together, these findings imply that RDC-dependent cluster transcripts are required to recruit the full complement of ping-pong factors to nuage. However, nuclear export of these transcripts appears to be mediated by TREX-dependent and TREX independent mechanisms, and TREX-independent export is sufficient to support Vasa and Ago3 localization to nuage and low levels of piRNA production in *mnk, uap56* and mnk, *thoc7* mutants. By contrast, Armi fails to localize in *rhi,* uap56 or *thoc7* mutants, and localization is not restored in the *mnk* double mutants. TREX-dependent export of piRNA precursor transcripts thus appears to be required for Armi localization to nuage.

The Armi helicase localizes to nuage and mitochondria and functions with Zuc in phased biogenesis of transcripts produced by Ago3, suggesting that Armi shuttles precursors between nuage and mitochondria (Ge et al., 2019). However, RDC and TREX mutants and the corresponding *mnk* double mutants displace Armi from nuage, but do not disrupt phased piRNA biogenesis (Figure 4C and S6C). Armi accumulation at nuage thus appears to be dispensable for phasing. However, Armi could rapidly cycle in and out of nuage, and the RDC and TREX mutations could decrease the rate of Armi association with nuage, or increase the rate of Armi exit, thus reducing steady state accumulation without blocking transient association with this structure. The functional significance of Armi accumulation at nuage thus remains to be definitively established.

### TREX function in spatial juxtaposition of nuage and clusters

In wild type ovaries, prominent nuage granules localize to regions of the nuclear periphery opposite piRNA clusters. *uap56* and *thoc7* single mutations disrupt localization of most nuage components, leading us to propose that TREX-mediated export of cluster transcripts promotes nuage assembly. However, our studies show genetically perturbating of *mnk* restores nuage localization of Aub, Ago3, and Vasa in *uap56* mutants. Thus, revealing activation of Chk2 disrupts nuage composition and prompting us to reevaluate nuage position relative to clusters in *uap56* and *mnk, uap56* mutants, and in *w^1^* controls. As shown in Figure 5, quantification of nuage localization relative to cluster indicates that the nuage granules that form in the double mutants are not biased toward regions of the nuclear envelope opposite clusters. Therefore, TREX-dependent precursor export from the nucleus biases the location of perinuclear nuage.

### Nuage assembly

Phase separated granules are proposed to have critical functions in numerous nuclear and cytoplasmic processes, including piRNA biogenesis. *Drosophila* Vasa, the first molecularly characterized nuage component, is required for ping-pong amplification, and is reported to have an essential function in nuage assembly (Liang et al., 1994; Xiol et al., 2014). However, we find that Ago3 and Armi localizes to perinuclear granules in *vas* null mutant, and observe electron dense perinuclear nuage in two different *vas* null mutant combinations (Figure 3, S3C and S3D). In addition, thin section EM revealed electron dense perinuclear nuage in *aub, ago3, armi, rhi, uap56* and *thoc7* mutant backgrounds (Figure 3, S3, S4). Nuage assembly thus does not require any of the individual products of these genes, including Vasa.

While none of the mutations analyzed here disrupt nuage assembly, they define a genetic hierarchy controlling nuage composition (Figure 7). As noted above, *rhi* mutations, which disrupt RDC-dependent cluster transcription, displace Armi, Aub, Ago3 and Vasa localization to perinuclear nuage, and localization is not restored in *mnk* double mutants. Cluster transcripts thus appear to recruit the ping-pong machinery and Armi to nuage. Within nuage, we find that Aub localization requires both Vasa and Ago3, but Ago3 and Vasa localize independent of each other, and independent of Aub. Consistent with an upstream function for Ago3 in the phased biogenesis pathway, *ago3* mutations block Armi localization to nuage, and *armi* mutations to do not alter nuage localization of Ago3 or other nuage components (Figure S3). These observations support a model in which RDC-dependent transcripts recruit Ago3 and Vasa to nuage, and Aub is localized to nuage through direct or indirect interactions with Vasa and Ago3. Within this framework, we speculate that Chk2 activation targets Vasa, displacing both Vasa and Aub (Figure 7). piRNA protein localization studies suggest that piRNA biogenesis proceeds though a series of tightly linked nuclear, perinuclear and mitochondrial compartments. However, our analysis of a novel *vas* allele indicating that disrupting nuage, the central component of this system, has minimal impact on piRNA biogenesis and transposon silencing.

**Figure 7:**
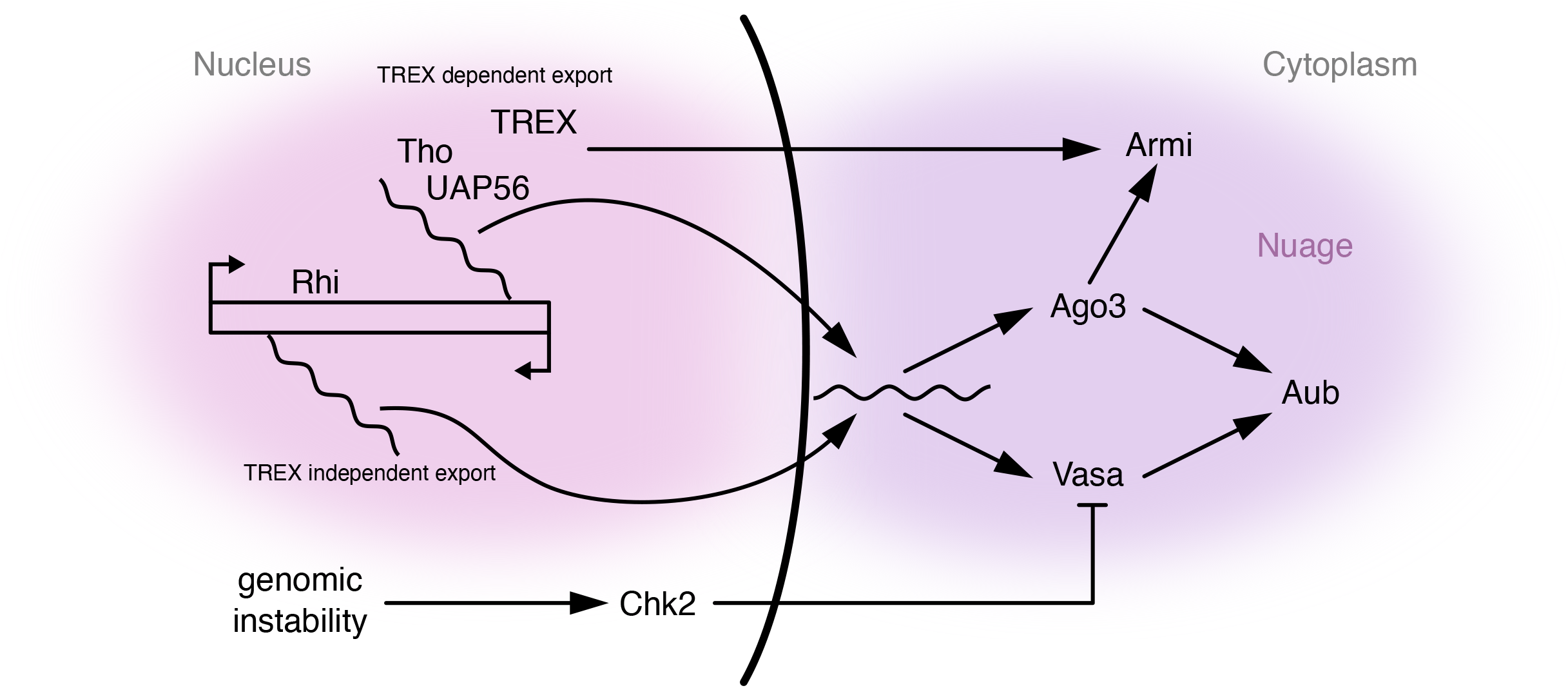
Model of Chk2 modulating nuage composition and interdependency between piRNA pathway proteins and cluster transcripts. Rhino bound to dual stranded piRNA clusters initiate transcription of long piRNA precursors from both genomic strands. We propose that that these transcripts can be exported out of the nucleus in a TREX dependent and TREX independent manner. piRNA precursors the cytoplasm can facilitate Ago3 and Vasa localization to nuage granules and their localization is independent from each other. However, Aub localization to nuage is dependent on Ago3 and Vasa. Lastly Armi localization to nuage requires intact TREX complex and Ago3. Upon genomic instability, activation of DNA damage signaling though Chk2 affect Vasa and Aub localization to nuage. We propose Chk2 signaling may target Vasa and Aub displacement may be a downstream effect loss of Vasa at nuage.

### Nuage function

Biochemical studies indicate that Vasa promotes release of cleaved products from PIWI proteins during the ping-pong cycle (Xiol et al., 2014), and null *vas* mutations have a profound impact on ping-pong amplification and transposon silencing (Figure S6 and S7). Vasa also localizes to the posterior pole of the oocyte, and these mutations disrupt posterior patterning and germline specification during embryogenesis. The *vas^PD^* allele was previously shown to disrupts pole plasm assembly and produce very low levels of protein that is only detected within the germarium, during the earliest stages of oogenesis (Johnstone & Lasko, 2004; Lasko & Ashburner, 1990). Our RNAseq analysis indicates that this allele produces wild type protein, but at levels approximately 100 fold lower than wild type. We also find that Aub, which is the primary driver of transposon transcript cleavage and post-transcriptional silencing, fails to localized to nuage in *vas^PD^* mutants (Figure 6B). However, piRNA expression, ping-pong bias and transposon silencing are comparable to wild type controls. Localization of Aub, and continued expression of Vasa protein, thus appear to be dispensable for piRNA amplification and transposon silencing through most of oogenesis. Furthermore, these findings suggest that Vasa is required to initiate the ping-pong amplification, but is not required to propagate the cycle, which is presumably sustained by Aub and Ago3. In addition, we find that phased piRNA production is unaltered in *rhi, uap56* and *thoc7* mutants, which displace Armi from nuage.

The findings reported here thus link perinuclear nuage assembly to Rhino-dependent piRNA precursor assembly, define interactions between the cytoplasmic ping-pong proteins that control nuage composition, and reveal a novel function for Chk2 damage signaling in control of nuage organization and function. However, these findings also indicate that nuage localization of Aub, Vasa or Armi is dispensable for ping-pong amplification, phased piRNA biogenesis and transposon silencing, implying that these critical processes take place in the cytoplasm, not in nuage. The function of these highly conserved phase separated structures thus remains to be determined. However, Aub binds anti-sense piRNAs and drives post-transcriptional transposon silencing. We observed that Aub is liberated from nuage by Chk2 signing and this raises the possibility that nuage could therefore buffers transposon silencing activity by releasing effectors when transposons are mobilized and Chk2 is activated, which in turn helps maintain germline genome integrity.

## Acknowledgements

We would like to thank current and past members of the Theurkauf and Weng labs for their insightful discussion and comments throughout the project; Socheata Ly, Silvia Corvera, and David Grunwald for insightful discussions on image analysis; Ildar Gainetdinov for sharing bioinformatic analysis looking for phased piRNA biogenesis; Jean-René Huynh for *rpp30* stocks; Keith Reddig and the Umass Chan EM core facility for Electron microscopy services. This work was supported by NIH grants R01-HD049116 to W.T.

## STAR methods

### Key Resources Table

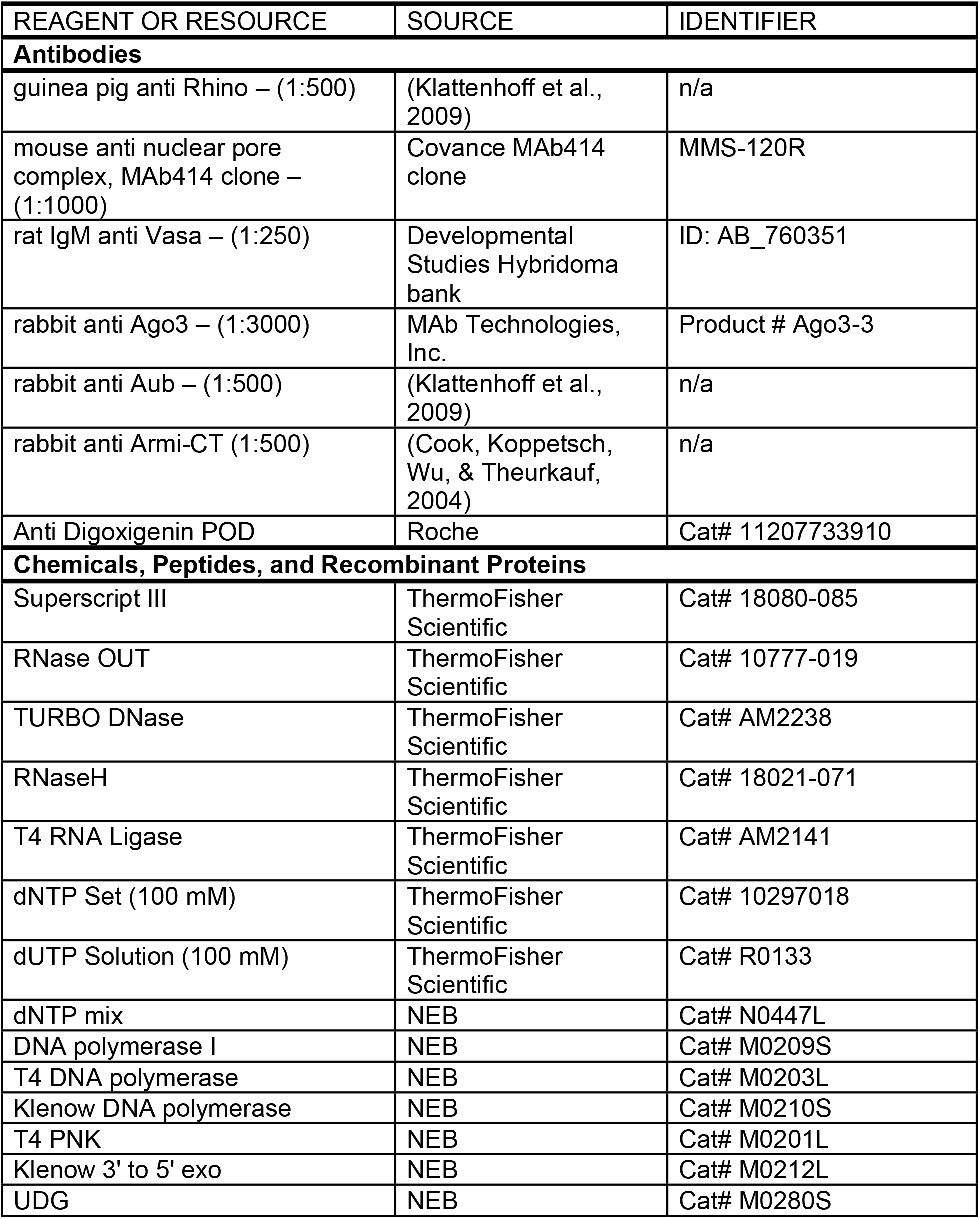
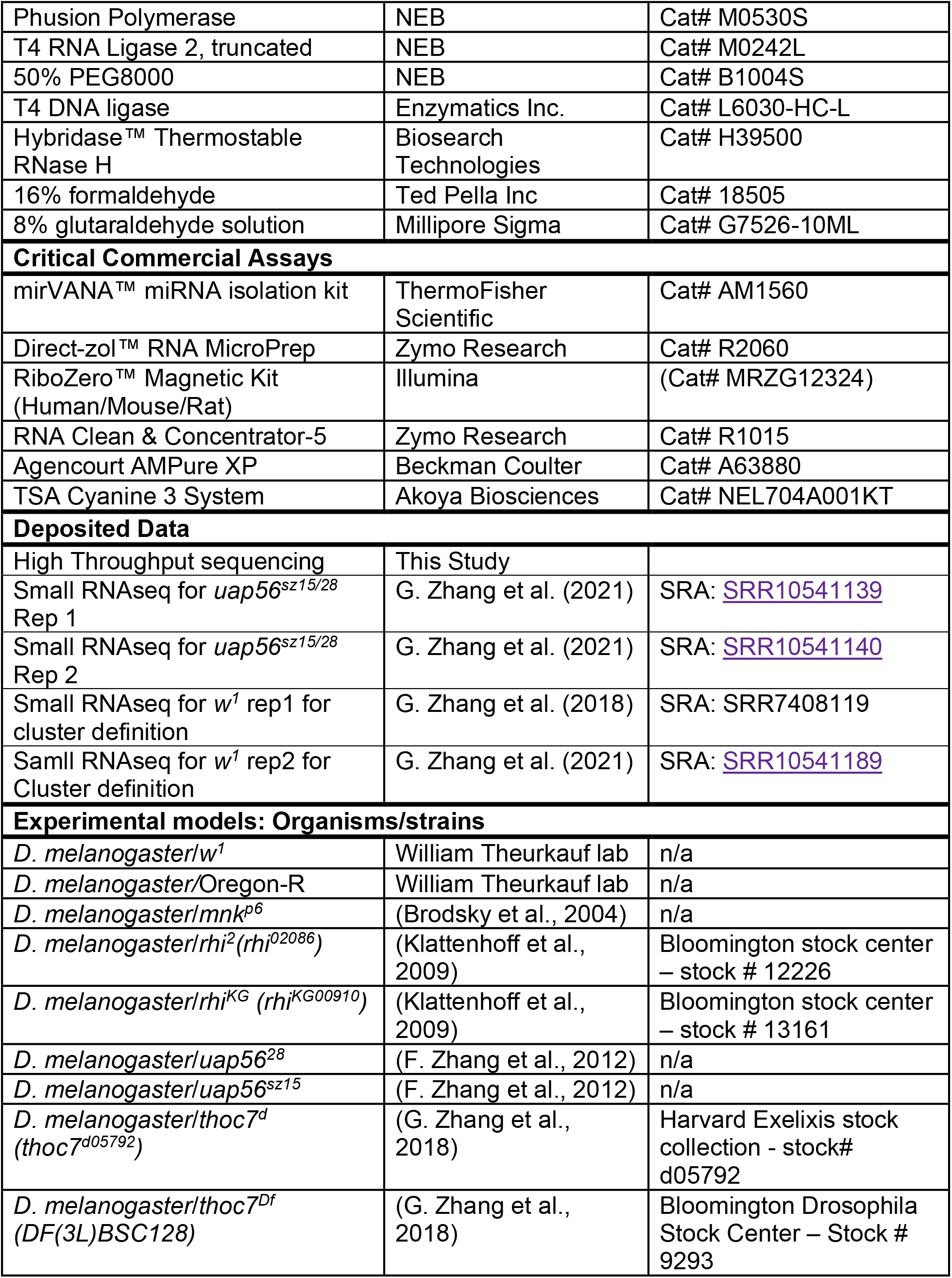
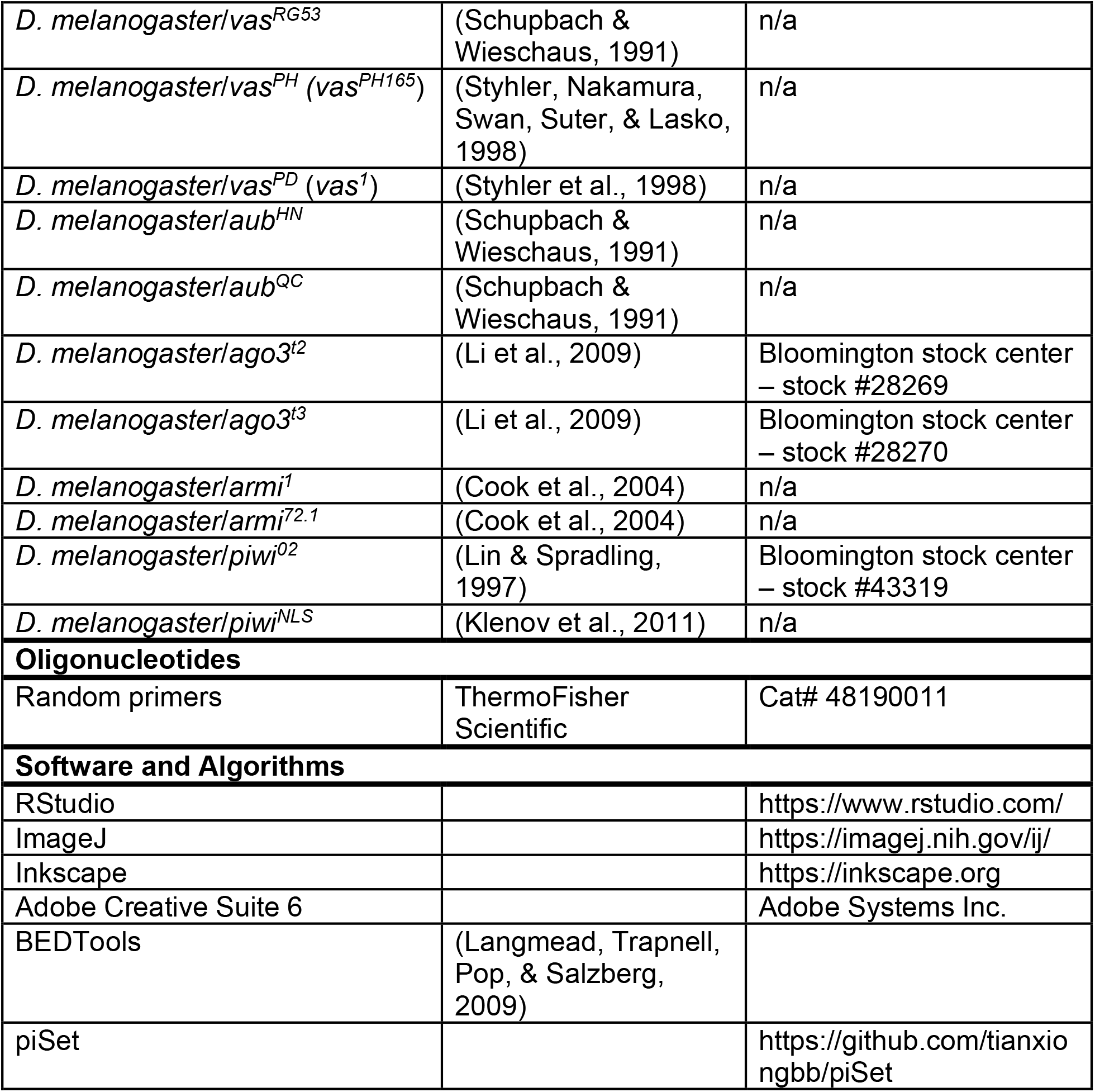

### Fly strains and husbandry

All fly crosses were maintained at 25°C on cornmeal medium except for crosses generating *thoc7* and *mnk; thoc7* mutants which were raised between 18°C-20°C. *mnk; thoc7^d/Df^* double mutants were not recoverable with crosses maintained at 25°C. Females were collected at 1-2 day old and fed on yeast for 2 days. Ovaries were collected from 2-4 day old females unless otherwise stated. Standard genetic procedures were used to generate double mutant combinations and generating mutant trans-hets.

### Immunofluorescence

Protocol previously described in (McKim, Joyce, & Jang, 2009) and F. Zhang et al., 2012. Briefly, 2-4 day old female ovaries were dissected in Robb’s media, fixed in 4% formaldehyde, washed, incubated in primary overnight, washed, incubated in fluorophore conjugated secondary and DAPI for DNA staining overnight, washed, and mounted on slide with mounting medium.

## Image analysis

### Automated Quantification of perinuclear nuage localization

z-stacks were taken of nurse cell nuclei stained for two different nuage markers and nuclear pore. Images were analyzed in ImageJ, where signal was defined as four standard deviations above the mean for each marker. Signal for the two nuage markers were merged and used to define nuage particles that were larger than 0.013-0.026 μm^2^ in size. Only particles within 0.25-0.36 μm proximity to the nuclear membrane signal were considered perinuclear granules. Signal intensity of each nuage marker was then measured for every granule identified previously. See extended methods for annotated scripts used.

### Automated quantification of nuclear and cytoplasmic spatial juxtaposition

To measure the percent of peripheral Rhino juxtaposed with the cytoplasmic nuage in an unbias manner, we developed an automated image analysis pipeline using ImageJ. z-stacks were taken of nuclei stained for Rhino, nuclear membrane, and nuage marker. Signal was defined as 3 standard deviations above the mean signal for each marker. The nuclear membrane signal was used to identify any Rhino and nuage proximal to the nuclear membrane. Total number of Rhino at the periphery was counted. To count how many peripheral Rhino was adjacent to nuage marker, we looked for overlap between nuage and rhino signal. Normally, nuage and Rhino signal do not colocalize, therefore, we expanded the nuage marker signal equally in all directions and measured overlap with each exaptation. A range of 0-6 pixel expansion were tested. All expansions showed similar trends as to each other and to the manual method of measuring spatial juxtaposition described in F. Zhang et al., 2012. Expansion 3 was chosen moving forward. See extended methods for annotated scripts used.

### Electron microscopy

Ovaries were collected from 2-4 day old Females and fixed overnight in fixative buffer containing 2% glutaraldehyde and 100 mM sodium cacodylate. Dehydration, embedding, sectioning, and staining were done at the Umass Chan Medical School EM core facility.

### Ovary Small RNA seq

Small RNA-seq library construction was described previously (G. Zhang et al., 2021). Briefly, total RNA was extracted from 2-4 days old female ovaries using the MirVana kit (Ambion). Small RNAs 18-29 nucleotide long were isolated using polyacrylamide gel purification. Sequencing libraries were made by first depleting the 2S rRNA followed by 3ʹ adaptor ligation, gel purification, 5ʹ adaptor ligation, gel purification, Reverse transcription, and PCR amplification. Libraries were sequenced using Illumina Nextseq. In general, two or more biological replicates was generated for each genotype.

### Ovary Long RNA seq

Strand specific long RNA-seq library construction was described previously (G. Zhang et al., 2021). Briefly, total RNA was extracted from 2-4 days old female ovaries using the MirVana kit (Ambion). rRNA was depleted using antisense rRNA oligo hybridization followed by RNase H digestion. Library construction consisted of RNA Fragmentation, Reverse Transcription, dUTP incorporation, end repair, size selection using Ampure XP beads, A-tailing, Adaptor ligation, UDG treatment and PCR amplification. Libraries were sequenced using the Illumina Nextseq. In general, two or more biological replicates was generated for each genotype.

### Bioinformatics analysis

The bioinformatic analysis was performed as described previously (G. Zhang et al., 2021). The *Drosophila* reference genome (*dm6*), rRNA sequences, gene annotations, and hairpin sequences are from Flybase (Version 6.13). Transposon consensus sequence were from Repbase (Bao, Kojima, & Kohany, 2015).

### piRNA cluster annotation

We used a similar method as Yu et al. (2019) to annotate piRNA clusters in the dm6 genome using *w^1^* control small RNA-seq data from G. Zhang et al. (2021) (See Key Resource table). We considered 24–32 nt small RNA reads that could map to the *dm6* genome, after rRNA, miRNA, tRNA, snRNA, and snoRNA were removed, as piRNAs. piRNAs were then assigned to 20 kb sliding windows (with a 1 kb step), and windows with more than 100 piRNAs per million uniquely mapped piRNAs were further considered as potential piRNA clusters. To remove false positives due to un-annotated miRNA, rRNA, tRNA, snRNA, and snoRNA, which mostly produce reads with the same sequences, we also filtered out those 20-kb genomic windows with fewer than 200 distinct reads (called species). We then calculated the first-nucleotide content for each 20-kb window, and those windows with 1U/10A percentage less than 50% were also discarded. The remaining contiguous 20-kb windows were deemed putative piRNA clusters. Finally, we performed manual curation for putative piRNA clusters using piRNA profile. Determined by the direction of the piRNAs produced, piRNA clusters are classified as uni-strand and dual-strand piRNA clusters.

### RNA-Seq Analysis

piSet_rnaseq pipeline from GitHub was used to analyze long RNA sequences reads. Briefly, Bowtie2 (Version 2.2.5) with default settings was used to map raw sequencing reads to rRNA sequence (Langmead & Salzberg, 2012). The remaining reads were mapped to the *Drosophila* genome (*dm6*) and transposon consensus sequences using STAR (Version 2.5.2b) and Hisat2 with default parameters (Dobin et al., 2013; Kim, Langmead, & Salzberg, 2015). The transcript abundance for each gene, transposon, and cluster (RPKM: Reads Per Kilobase per Million mapped reads) was counted using BEDTools (Version 2.27.1) (Quinlan & Hall, 2010) and normalized to total number of genome mapping reads after excluding rRNA mapping reads.

### Small RNA-seq Analysis

piSet_srnaseq was used to analyze small RNA sequencing reads. Briefly, the 3ʹ end adaptor was removed via cutadapt (Version 1.15) (Martin, 2011) and the raw small RNA-seq reads were mapped to rRNA, miRNA hairpin, snoRNA, snRNA, and tRNA sequence allowing for no mismatches using Bowtie (Version 1.1.0) (Langmead et al., 2009). The remaining reads were mapped to the *Drosophila* genome (*dm6*) and transposon consensus sequences. Small RNA abundance was normalized to reads mapping to *Flamenco*. For Ping-Pong analysis, the 5ʹ go 5ʹ overlaps between piRNAs mapping to opposite genomic strands were calculated, and Z-score for 10 nucleotide (nt) overlap was calculated by using 1-9 nt and 11-30 nt overlaps as background (Li et al., 2009). For Phasing analysis, the shortest distance between the 3ʹ end of piRNAs and the 5ʹ end of the piRNA downstream was measured. Z-score for 0 nt between piRNAs was calculated using 1-19 nt as background (Han et al., 2015).

