## Supplemental Figures for "Aub, Vasa and Armi localization to phase separated nuage is dispensable for piRNA biogenesis and transposon silencing in *Drosophila*"

#### Supplemental Figure Legends

**Figure S1: Additional nuage colocalization conditions on wildtype *w*$^1$**

A-C. Immunofluorescence for two nuage markers Vasa (green) and Armi (red) in *w*$^1$ show prominent Vasa containing granules colocalize with Armi (filled arrow heads). Some Vasa containing granules lack Armi (open arrow heads). Scale bar is 2 μm.

D-F. Immunofluorescence for two nuage markers for Ago3 (green) and Aub (red) in *w*$^1$ show high levels of colocalization (filled arrow heads). Scale bar is 2 μm.

G. Density scatterplot comparing of average Ago3 signal vs average Aub signal where each point represents a nuage granule in *w*$^1$. Only granules proximity to nuclear pore signal was used for this analysis. Pearson correlation coefficient was calculated.

H. Violin plot showing distribution of Aub/Ago3 signal ratios.

**Figure S2: Ago3, Vasa colocalization and nuage composition *uap56* single and *mnk, uap56* double mutants**

A-I. Immunofluorescence for Vasa (green) and Ago3 (red), in heterozygous controls, *uap56*$^{sz15/28}$, and *mnk, uap56*$^{sz15/28}$ double mutants. Scale bar is 2 μm.

J-L. Density scatterplot comparing of average Vasa signal vs average Ago3 signal where each point represents a perinuclear nuage granule for heterozygous controls, *uap56*$^{sz15/28}$, and *mnk, uap56*$^{sz15/28}$ double mutants respectively. Pearson correlation coefficient was calculated for each genotype.

M. Immunofluorescence for Armi was perfumed in *w*$^1$controls, *uap56*$^{sz15/28}$, and *mnk, uap56*$^{sz15/28}$ double mutants. Armi localization is depleted from nuage in both *uap56*$^{sz15/28}$, and *mnk, uap56*$^{sz15/28}$ double mutants. Scale bar is 2 μm.

**Figure S3: Localization of Vasa, Aub, Ago3 and Armi in various piRNA single and *mnk* double mutants reveals dependencies on nuage organization**

A. Vasa immunofluorescence staining in WT ($w^{1}$) and various piRNA mutants. Each image is an optical z-section through a nurse cell nucleus.

B. Aub immunofluorescence staining in WT ($w^{1}$) and various piRNA mutants.

C. Ago3 immunofluorescence staining in WT ($w^{1}$) and various piRNA mutants.

D. Armi immunofluorescence staining in WT ($w^{1}$) and various piRNA mutants.

**Figure S4: nuage still present in other piRNA mutants**

A-D. Electron microscopy images of Stage 3-5 nurse cells for $rhi^{KG/02}$, $thoc7^{d/Df}$, $aub^{HN/QC}$, and *mnk,* $armi^{1/72.1}$ mutants respectively. Electron dense nuage structures designated by the black arrow heads still detected in all genotypes. N (nucleus), C (cytoplasm), and M (mitochondria).

**Figure S5: piRNA expression, transposon silencing, and cluster expression is comparable between $w^{1}$ and *mnk***

A. Scatter plot comparing $w^{1}$ control with *mnk* mutants for antisense transposon mapping piRNAs (log 10) and transposon ping-pong z-score (log 2). Each point is a transposon family. Red points indicate more than two-fold up and blue points show more than two-fold down.

B. Scatterplot comparing $w^{1}$ control with *mnk* mutants for transposon and cluster expression where each point represents a transposon family or piRNA cluster respectively. Red points indicate more than two-fold up and blue points show more than two-fold down.

**Figure S6: Transposon mapping piRNA comparisons between single piRNA mutants and *mnk* double mutants reveals Chk2 effect on ping-pong amplification**

A. Scatter plot showing antisense transposon mapping piRNA abundance (log10) for Pair wise comparisons of WT($w^{1}$), piRNA single mutant and mnk double mutants. The following piRNA mutants were tested: $rhi^{KG/02}$, $thoc7^{d/Df}$, $vas^{RG/PH}$, $aub^{HN/QC}$, $ago3^{t2/t3}$, and $armi^{1/72.1}$. Red points indicate more than two-fold up and blue points show more than two-fold down.

B. Scatter plots showing ping pong z-score (log2) for each transposon family with the same comparisons as above.

C. Graphs showing the distribution of the shortest distance between 3' end of one piRNA and the 5' end of the subsequent piRNA downstream for various piRNA single mutants and mnk double mutants. Prominent peak at zero nucleotide is detected for all genotypes tested signifying intact phased piRNA biogenesis.

**Figure S7: Comparisons between single piRNA mutants and *mnk* double mutants for transposon and piRNA cluster expression show a subset of transposons are sensitive to DNA damage signaling**

A. Scatter plot showing pairwise comparisons of transposon expression levels between WT ($w^{1}$), single piRNA mutants and *mnk* double mutants. The following piRNA mutants were tested: *rhi*$^{KG/02}$, *thoc7*$^{d/Df}$, *vas*$^{RG/PH}$, *aub*$^{HN/QC}$, *ago3*$^{t2/t3}$, and *armi*$^{1/72.1}$. Red and blue points are more than two-fold up or down respectively.

B. Scatterplot showing piRNA cluster expression levels for the same comparisons as above. Each point is a piRNA cluster.

Figure S1

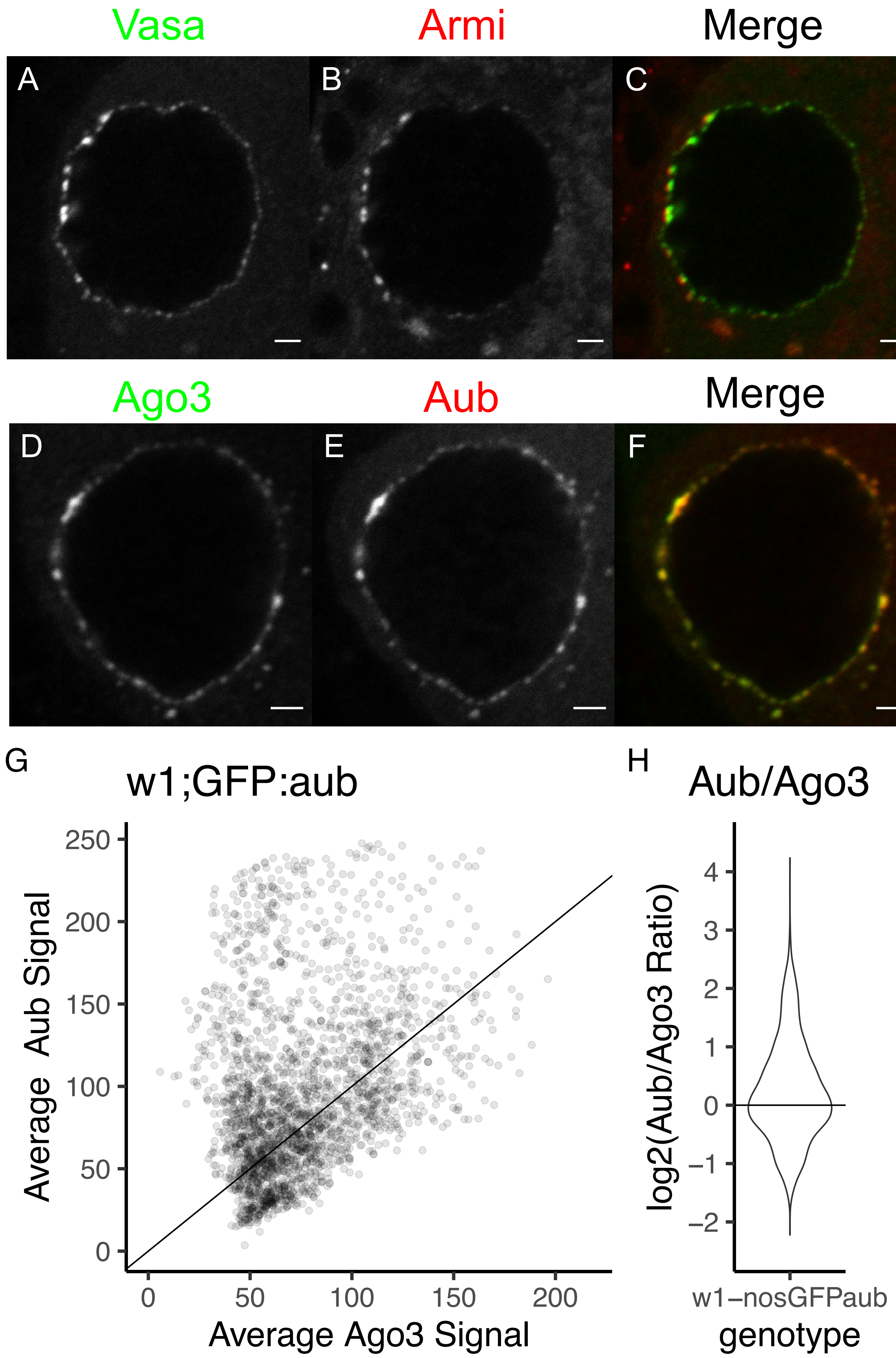
Vasa
Armi
Merge
A
B
C
Ago3
Aub
Merge
D
E
F
G
w1;GFP:aub
Average Aub Signal
Average Ago3 Signal
0
50
100
150
200
250
H
Aub/Ago3
log2(Aub/Ago3 Ratio)
−2
−1
0
1
2
3
4
w1−nosGFPaub
genotype

### Figure S2

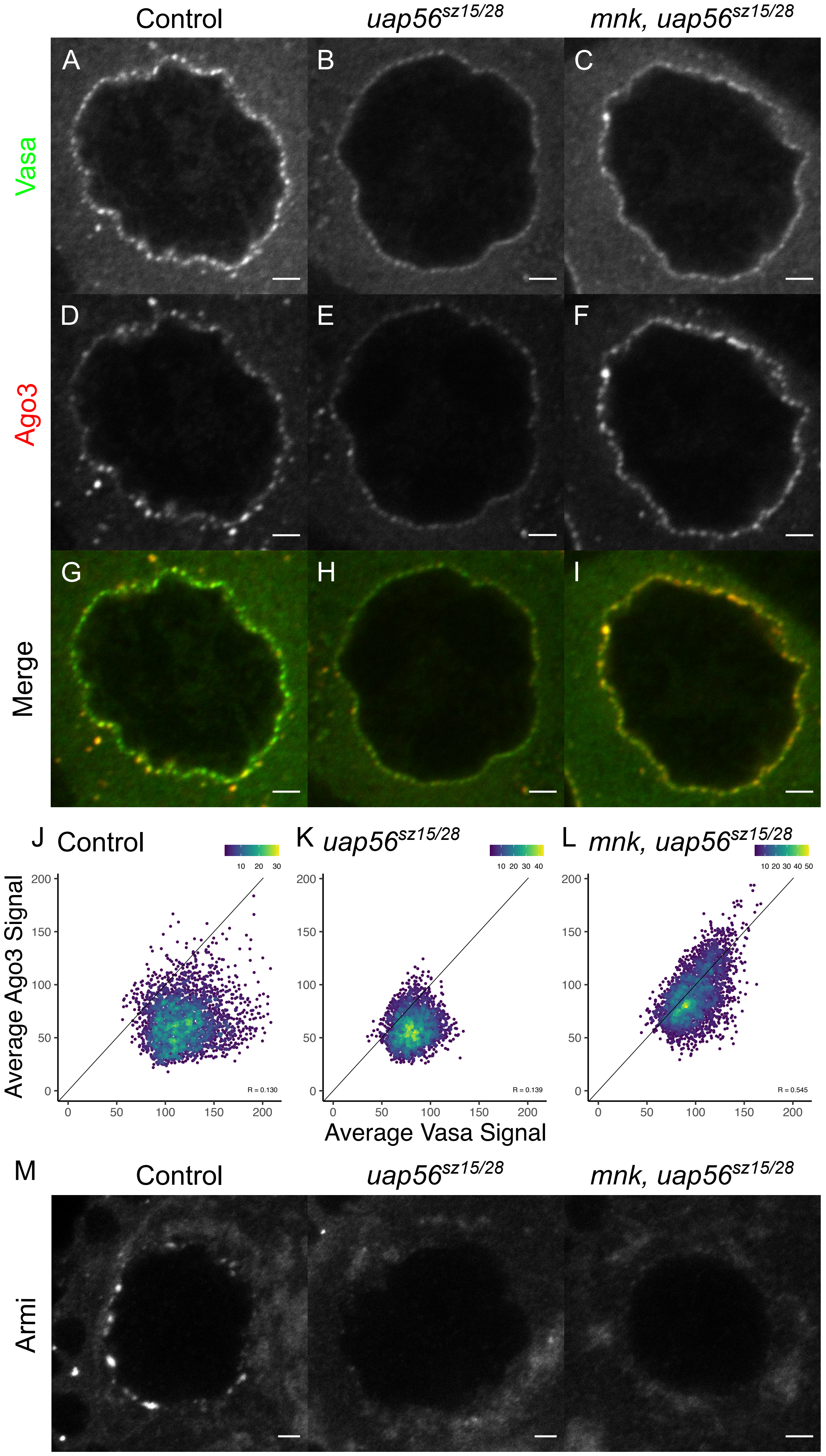

Control
uap56$^{sz15/28}$
mnk, uap56$^{sz15/28}$
A
B
C
Vasa
D
E
F
Ago3
G
H
I
Merge
J Control
K uap56$^{sz15/28}$
L mnk, uap56$^{sz15/28}$
Average Ago3 Signal
Average Vasa Signal
R = 0.130
R = 0.139
R = 0.545
M
Control
uap56$^{sz15/28}$
mnk, uap56$^{sz15/28}$
Armi

### Figure S3

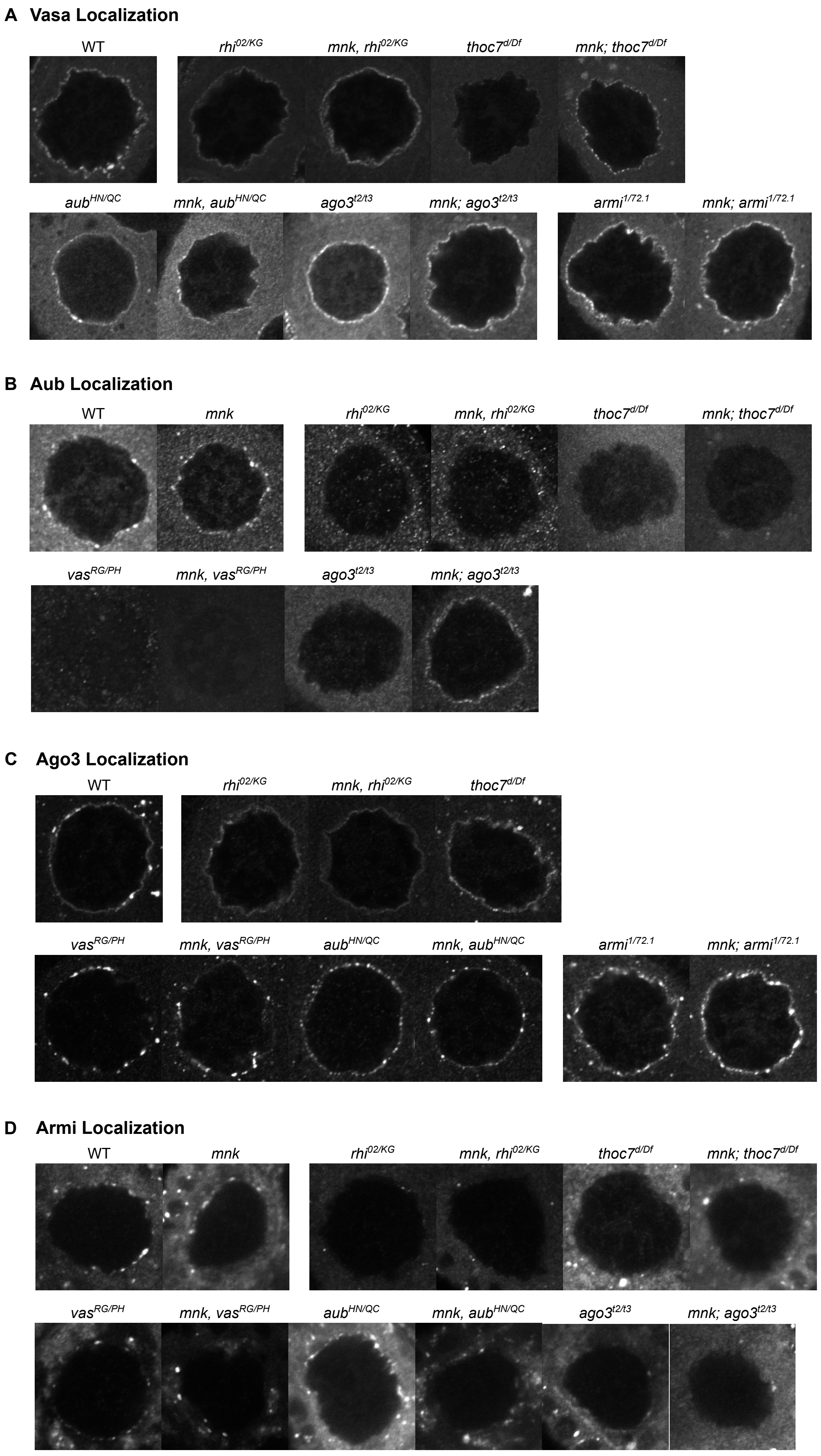
A Vasa Localization
WT
*rhi*$^{02/KG}$
*mnk, rhi*$^{02/KG}$
*thoc7*$^{d/Df}$
*mnk; thoc7*$^{d/Df}$
*aub*$^{HN/QC}$
*mnk, aub*$^{HN/QC}$
*ago3*$^{t2/t3}$
*mnk; ago3*$^{t2/t3}$
*armi*$^{1/72.1}$
*mnk; armi*$^{1/72.1}$
B Aub Localization
WT
*mnk*
*rhi*$^{02/KG}$
*mnk, rhi*$^{02/KG}$
*thoc7*$^{d/Df}$
*mnk; thoc7*$^{d/Df}$
*vas*$^{RG/PH}$
*mnk, vas*$^{RG/PH}$
*ago3*$^{t2/t3}$
*mnk; ago3*$^{t2/t3}$
C Ago3 Localization
WT
*rhi*$^{02/KG}$
*mnk, rhi*$^{02/KG}$
*thoc7*$^{d/Df}$
*vas*$^{RG/PH}$
*mnk, vas*$^{RG/PH}$
*aub*$^{HN/QC}$
*mnk, aub*$^{HN/QC}$
*armi*$^{1/72.1}$
*mnk; armi*$^{1/72.1}$
D Armi Localization
WT
*mnk*
*rhi*$^{02/KG}$
*mnk, rhi*$^{02/KG}$
*thoc7*$^{d/Df}$
*mnk; thoc7*$^{d/Df}$
*vas*$^{RG/PH}$
*mnk, vas*$^{RG/PH}$
*aub*$^{HN/QC}$
*mnk, aub*$^{HN/QC}$
*ago3*$^{t2/t3}$
*mnk; ago3*$^{t2/t3}$

**Figure S4**

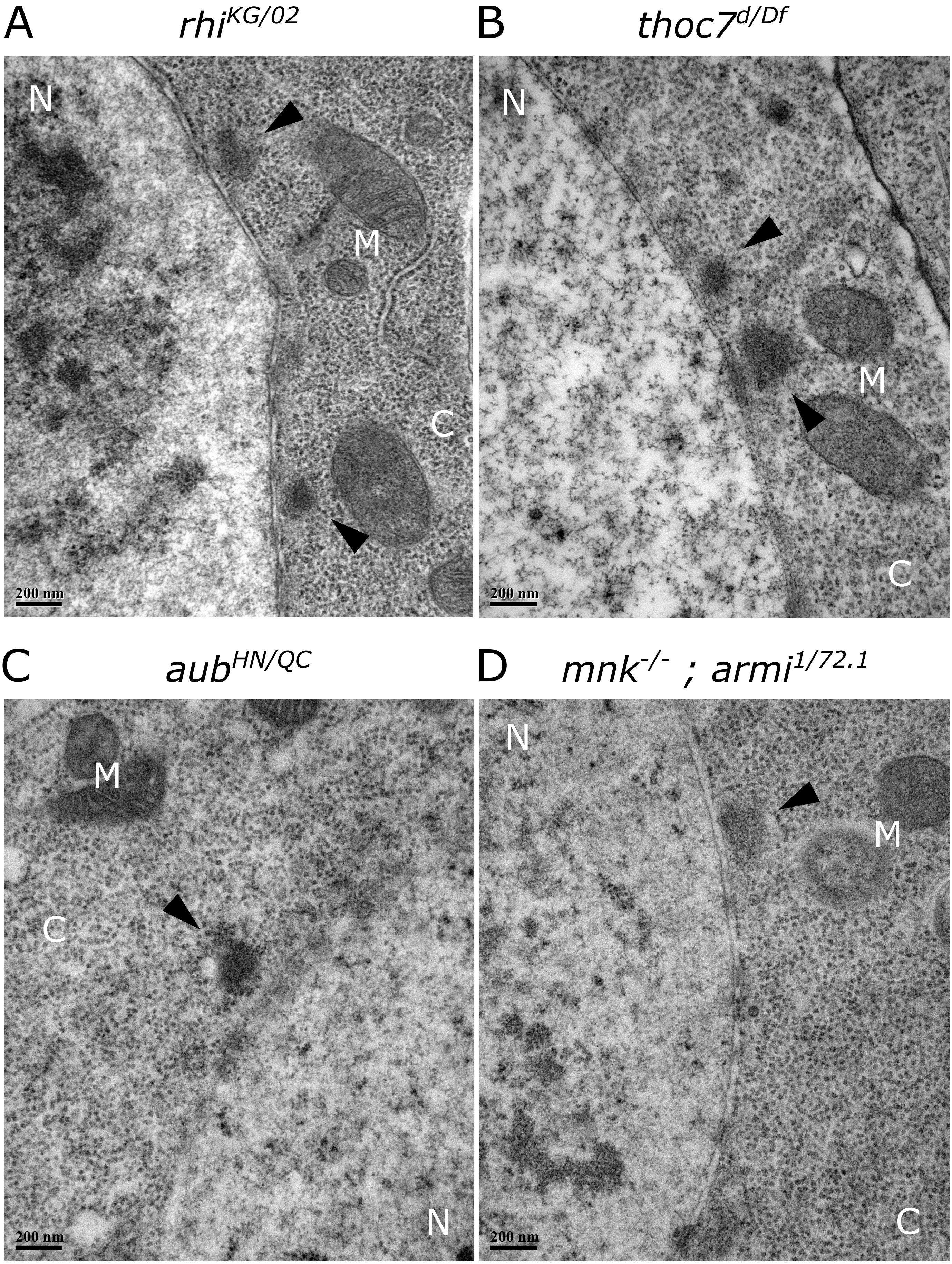
A
rhi^KG/02
N
M
C
200 nm
B
thoc7^d/Df
N
M
C
200 nm
C
aub^HN/QC
M
C
N
200 nm
D
mnk^-/- ; armi^1/72.1
N
M
C
200 nm

**Figure S5**

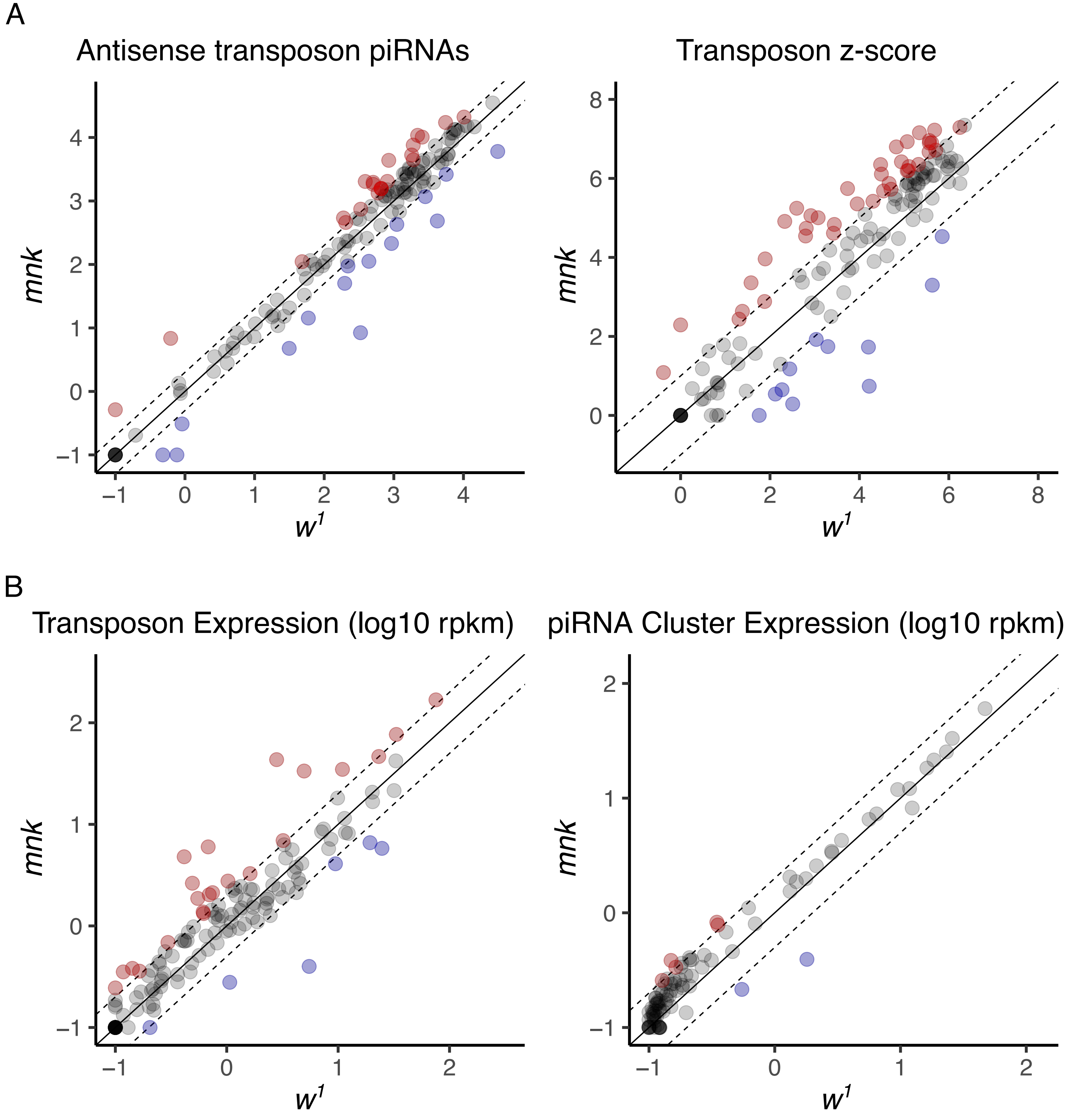
A
Antisense transposon piRNAs
Transposon z-score
B
Transposon Expression (log10 rpkm)
piRNA Cluster Expression (log10 rpkm)
mnk
w1

### Figure S6

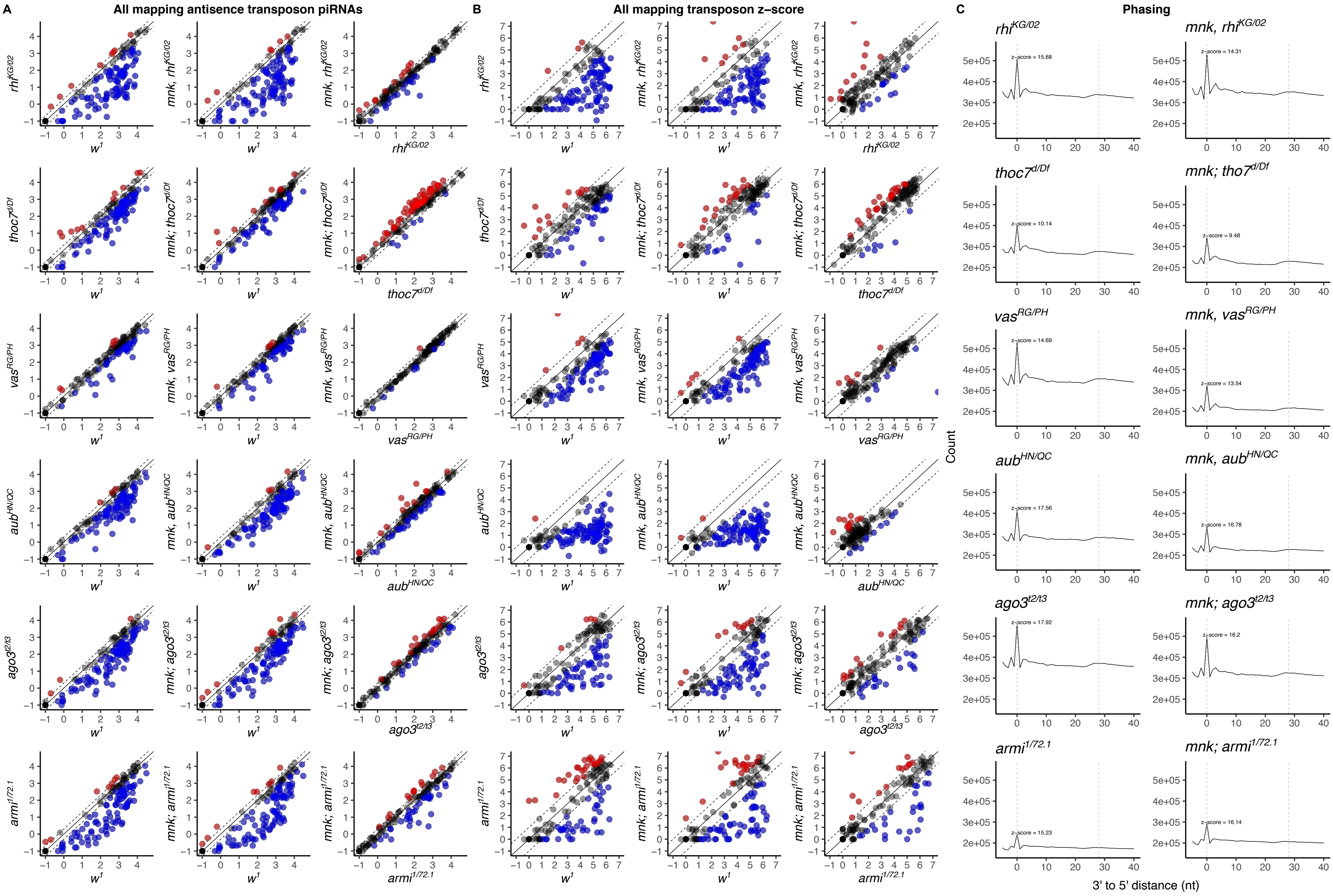

A
All mapping antisence transposon piRNAs
B
All mapping transposon z-score
C
Phasing
rhi^KG/02
mnk, rhi^KG/02
thoc7^d/Df
mnk; thoc7^d/Df
mnk; tho7^d/Df
vas^RG/PH
mnk, vas^RG/PH
aub^HN/QC
mnk, aub^HN/QC
ago3^t2/t3
mnk; ago3^t2/t3
armi^1/72.1
mnk; armi^1/72.1
w^1
Count
3' to 5' distance (nt)
z-score = 15.68
z-score = 14.31
z-score = 10.14
z-score = 9.48
z-score = 14.69
z-score = 13.54
z-score = 17.56
z-score = 16.78
z-score = 17.92
z-score = 16.2
z-score = 15.23
z-score = 16.14

### Figure S7

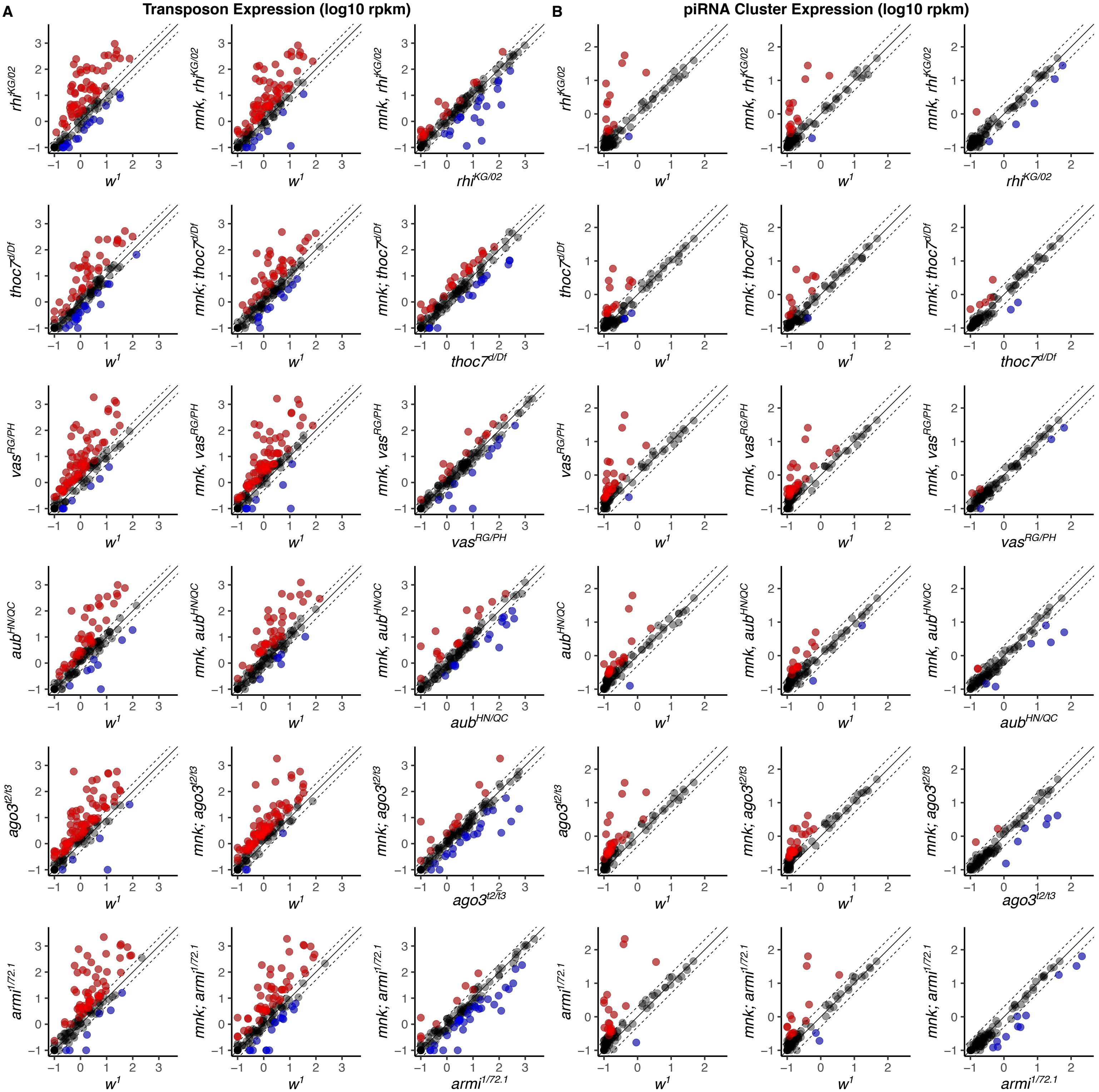
A
Transposon Expression (log10 rpkm)
B
piRNA Cluster Expression (log10 rpkm)
rhi^KG/02
mnk, rhi^KG/02
thoc7^d/Df
mnk; thoc7^d/Df
vas^RG/PH
mnk, vas^RG/PH
aub^HN/QC
mnk, aub^HN/QC
ago3^t2/t3
mnk; ago3^t2/t3
armi^1/72.1
mnk; armi^1/72.1
w^1

#### Extended Methods

#### Quantifying perinuclear nuage localization script

```
/*
******* NOTES and READ ME *********
* Language: IJ1 Macro
Things to check/change before running script.
1. Enter full path for output directory in Section 1
2. Change your channel extension (i.e. "_ch00.tif", "_ch01.tif", etc.) and
       nuage marker name so the labeled markers are designated properly
       in the section 4
3. Change the Date of the experiment in section 4
4. May need to change stringency of thresholding depending on the markers used
       in sections 5 and 6. Currently threshold is defined as mean + 4*std but we
       have tested mean + 3*std and mean + 5*std.  Recommend testing various threshold
       conditions to see which best represents the signal observed in the raw data.
5. In Section 5, nuclear periphery is defined by expanding/"Dilate" the nuclear
       membrane signal by a set number of pixels. For the magnification we used,
       expanding by 7 pixels was optimum to capture entire nuage granule, but you
       may need to change this parameter depending on the magnification used.
6. In Section 8, when defining nuage particles, we required each particle to be larger
       than 10 pixels.  Again, depending on the magnification used, this parameter may
       be inappropriate.
7. file folder setup on Windows and Mac are designated a little differently and
       you may need to play with sections 1-4 to designate files appropriately

Input file setup:
       This script is designed to take the folder containing all the images for a
       single nucleus as the input. Be sure that the folder contains the merged image
       of all the labeled markers as well as each marker split out into its own channel.
       See below for an example where "prefix" is the input folder.  prefix_z01.tif,
       prefix_z02.tif, etc. are the merged images, and anything with _ch0#.tif are the
       split-out channels for each marker. The suffix of each file is how the scripts
       designates each marker, if your files suffixes differ, you can modify sections
       1-4 accordingly.

       prefix/
              prefix_z01.tif
              prefix_z01_ch00.tif
              prefix_z01_ch01.tif
              prefix_z01_ch02.tif
              prefix_z01_ch03.tif
              prefix_z02.tif
              prefix_z02_ch00.tif
              prefix_z02_ch01.tif
              prefix_z02_ch02.tif
              prefix_z02_ch03.tif
              prefix_z03.tif
              prefix_z03_ch00.tif
              ... etc.

Output files
       The average signal for each nuage marker can be found in the files ending with
       _perinuclear_results.csv.  Each row in the table represents a nuage granule
       defined in Section 8. (example: prefix_nuage1_perinuclear_results.csv and
       prefix_nuage2_perinuclear_results.csv)

       Images containing “threshold_peripheral” show the regions that passed thresholding
       for each nuage marker and the regions that are considered perinuclear.  Use these
       images to optimize conditions for thresholding and defining perinuclear granules.

       Images containing “Merge_peripheral” is the image that is used to define regions
       of interests.
*/


/* Section 1: pop-up asking for input directory and setting the output directory
*/
```

```
print("\\Clear");
input = getDirectory("Input directory");
output = "/Full_path_to/Output_directory";
print ("Input directory: " + input);
print ("Save directory: " + output);



/* Section 2: getting list of prefixes for each slice of the z stack
*/
allnames = getFileList(input)
// for (i=0; i<allnames.length; i++) print(allnames[i]);

for (i=0; i<lengthOf(allnames); i++) {
	if (endsWith(allnames[i], "ch00.tif")==false &&
	endsWith(allnames[i], "ch01.tif")==false &&
	endsWith(allnames[i], "ch02.tif")==false &&
	endsWith(allnames[i], "ch03.tif")==false) {
		print (allnames[i]);
		files = Array.concat(files, allnames[i]);
	}
}

files = Array.slice(files,1);
// for (i=0; i<prenames.length; i++) print(prenames[i])


/* Section 3: making directory for output with sorce image name
*/
Directory = replace(files[0],"_z00.tif","");
print(Directory)
File.makeDirectory(output+File.separator+Directory)


setBatchMode(true);

/* Section 4: defining prefixes for saving files and also different channels
for each marker stained
*/

for (i=0; i<lengthOf(files); i++) {
	nuage1 = replace(files[i],".tif","_ch01.tif");
	nuage1name = "vasa";
	nuage2 = replace(files[i],".tif","_ch03.tif");
	nuage2name = "ago3";
	nup = replace(files[i],".tif","_ch02.tif");
	prefix = replace(files[i],".tif","_");
	date = "20210110_";
	roiManager("reset");

/* Section 5: define nuclear membrane and expanding signal to define nuclear periphery
 */
	open(input+File.separator+nup);
		run("Subtract Background...", "rolling=50");
		getStatistics(area, mean, min, max, std);
		threshold = mean + 4*std;
		setThreshold(0, threshold);
		setOption("BlackBackground", false);
		run("Make Binary", "thresholded remaining black");
		run("Convert to Mask");
		run("Options...", "iterations=1 count=1 do=Nothing");
		run("Dilate");
		run("Dilate");
		run("Dilate");
		run("Dilate");
		run("Dilate");
		run("Dilate");
		run("Dilate");


/* Section 6: thresholding both nuage signal
```

```
 */
	open(input+File.separator+nuage1);
		run("Subtract Background...", "rolling=50");
		getStatistics(area, mean, min, max, std);
		threshold = mean + 4*std;
		setThreshold(0, threshold);
		setOption("BlackBackground", false);
		run("Make Binary", "thresholded remaining black");
		run("Convert to Mask");
	open(input+File.separator+nuage2);
		run("Subtract Background...", "rolling=50");
		getStatistics(area, mean, min, max, std);
		threshold = mean + 4*std;
		setThreshold(0, threshold);
		setOption("BlackBackground", false);
		run("Make Binary", "thresholded remaining black");
		run("Convert to Mask");


/* Section 7: creating images of just peripheral nuage markers
 */
	imageCalculator("AND create", nuage1, nup);
		selectWindow("Result of "+nuage1);
		saveAs("Tiff",
output+File.separator+Directory+File.separator+date+"threshold_peripheral_"+nuage1);
	imageCalculator("AND create", nuage2, nup);
		selectWindow("Result of "+nuage2);
		saveAs("Tiff",
output+File.separator+Directory+File.separator+date+"threshold_peripheral_"+nuage2);

/* Section 8: merging both peripheral nuage markers for selecting one set of ROIs
 */
	roiManager("reset");
	imageCalculator("Add create", date+"peripheral_raw_"+nuage1,
date+"peripheral_raw_"+nuage2);
		selectWindow("Result of "+date+"peripheral_raw_"+nuage1);
		saveAs("Tiff",
output+File.separator+Directory+File.separator+date+"_Merge_peripheral_"+prefix);
	run("Analyze Particles...", "size=10-Infinity pixel exclude summarize add");


/* Section 9: Measure signal of first and second nuage marker
 */
if (roiManager("count") >= 1){
	run("Set Measurements...", "area mean standard min integrated median redirect=None
decimal=3");

	open(input+File.separator+nuage2);
	roiManager("Measure");
	close();
	selectWindow ("Results");
	saveAs("Results",
output+File.separator+Directory+File.separator+date+prefix+nuage2name+"_perinuclear_results.csv")
;
	run("Close");

	open(input+File.separator+nuage1);
	roiManager("Measure");
	close();
	selectWindow ("Results");
	saveAs("Results",
output+File.separator+Directory+File.separator+date+prefix+nuage1name+"_perinuclear_results.csv")
;
	run("Close");
}

}

setBatchMode(false)
```

#### Quantifying Nuclear and cytoplasmic spatial juxtaposition

```
/*
******* NOTES and READ ME *********
* Language: IJ1 Macro
Things to check/change before running script.
1. Enter full path for output directory in Section 1
2. Change your channel extension (i.e. "_ch00.tif", "_ch01.tif", etc.) to designate
       Rhino, nuclear pore and nuage appropriately in the section 4
3. Change the Date of the experiment in section 4
4. May need to change stringency of thresholding depending on the markers used
       in sections 5 and 6. Currently threshold is defined as mean + 3*std but we
       have tested mean + 4*std and mean + 5*std.  Recommend testing various threshold
       conditions to see which best represents the signal observed in the raw data.
5. In Section 5, nuclear periphery is defined by expanding/"Dilate" the nuclear
       membrane signal by a set number of pixels. For the magnification we used,
       expanding by 3 pixels was optimum to capture foci directly adjacent to the
       nuclear pore signal but you may need to change this parameter depending on
       the magnification used.
6. In Section 8, when defining Rhino particles, we required each particle to be larger
       than 5 pixels.  Again, depending on the magnification used, this parameter may
       be inappropriate.
7. file folder setup on Windows and Mac are designated a little differently and
       you may need to play with sections 1-4 to designate files appropriately

Input file setup:
       This script is designed to take the folder containing all the images for a
       single nucleus as the input. Be sure that the folder contains the merged image
       of all the labeled markers as well as each marker split out into its own channel.
       See below for an example where "prefix" is the input folder.  prefix_z01.tif,
       prefix_z02.tif, etc. are the merged images, and anything with _ch0#.tif are the
       split-out channels for each marker. The suffix of each file is how the scripts
       designates each marker, if your files suffixes differ, you can modify sections
       1-4 accordingly.

       prefix/
              prefix_z01.tif
              prefix_z01_ch00.tif
              prefix_z01_ch01.tif
              prefix_z01_ch02.tif
              prefix_z01_ch03.tif
              prefix_z02.tif
              prefix_z02_ch00.tif
              prefix_z02_ch01.tif
              prefix_z02_ch02.tif
              prefix_z02_ch03.tif
              prefix_z03.tif
              prefix_z03_ch00.tif
              ... etc.

Output files
       Counting Rhino overlap with Nuage is found in "results_ex#.csv" files.  The
       each row represents a single Rhino focus or Region of Interest (RIO). Max
       column designates whether there is nuage overlap with that RIO.
              0 means no overlap and no adjacent localization of nuage
              255 means there is overlap and nuage is adjacent to Rhino Focus
       Percent of Rhino spatially juxtaposed to nuage can be calculated by counting
       the 0s and 255s. Adjacent localization does usually does not result in overlap,
       therefore, nuage signal is expanded equally and the overlap with RIOs are
       recorded with each expansion. ex0 is no expansion, ex1 is one pixel expansion,
       ex2 is two pixle expansion, etc.

       Files containing "summary.csv" are the output of particle picker.  The first row
```

```
        contains the number of Rhino added to the RIOs and every subsequent row measures
        the number of nuage particles with each expansion.

        Images containing “peripheral_raw” show the regions that passed thresholding
        and are considered perinuclear.  the different channels for each z-section of
        these images are merged to create the files containing “composite_raw”. All of
        these images can be used to optimize conditions for thresholding and defining
        perinuclear ROIs

        Images containing "composite_ex#.jpg" are the merged images as the nuage signal
        is expanded.  This can be used for visual inspection to determine which number
        of expansions is appropriate. ex0 is no expansion, ex1 is one pixel expansion,
        ex2 is two-pixel exaptation, etc.
         */




/* Section 1: pop-up asking for input directory and setting the output directory
*/
print("\\Clear");
input = getDirectory("Input directory");
output = "/Full_path_to/Output_directory";
print ("Input directory: " + input);
print ("Save directory: " + output);



/* Section 2: getting list of prefixes for each slice of the z stack
*/
allnames = getFileList(input)
// for (i=0; i<allnames.length; i++) print(allnames[i]);

for (i=0; i<lengthOf(allnames); i++) {
        if (endsWith(allnames[i], "ch00.tif")==false &&
        endsWith(allnames[i], "ch01.tif")==false &&
        endsWith(allnames[i], "ch02.tif")==false &&
        endsWith(allnames[i], "ch03.tif")==false) {
                print (allnames[i]);
                files = Array.concat(files, allnames[i]);
        }
}

files = Array.slice(files,1);
// for (i=0; i<prenames.length; i++) print(prenames[i])


/* Section 3: making directory for output with sorce image name
*/
Directory = replace(files[0],"_z00.tif","");
print(Directory)
File.makeDirectory(output+File.separator+Directory)


setBatchMode(true);

/* Section 4: defining prefixes for saving files and also different channels
for each marker stained
*/
for (i=0; i<lengthOf(files); i++) {
        nuage = replace(files[i],".tif","_ch02.tif");
        nup = replace(files[i],".tif","_ch00.tif");
        rhi = replace(files[i],".tif","_ch03.tif");
        prefix = replace(files[i],".tif","_");
        date = "20210911_";
        roiManager("reset");

/* Section 5: define nuclear membrane and dilate 3 times to define nuclear periphery
 */
        open(input+File.separator+nup);
                run("Subtract Background...", "rolling=50");
```

```
                getStatistics(area, mean, min, max, std);
                threshold = mean + 3*std;
                setThreshold(0, threshold);
                setOption("BlackBackground", false);
                run("Make Binary", "thresholded remaining black");
                run("Convert to Mask");
                run("Options...", "iterations=1 count=1 do=Nothing");
                run("Dilate");
                run("Dilate");
                run("Dilate");

/* Section 6: thresholding rhino and nuage signal
 */
        open(input+File.separator+rhi);
                run("Subtract Background...", "rolling=50");
                getStatistics(area, mean, min, max, std);
                threshold = mean + 3*std;
                setThreshold(0, threshold);
                setOption("BlackBackground", false);
                run("Make Binary", "thresholded remaining black");
                run("Convert to Mask");
        open(input+File.separator+nuage);
                run("Subtract Background...", "rolling=50");
                getStatistics(area, mean, min, max, std);
                threshold = mean + 3*std;
                setThreshold(0, threshold);
                setOption("BlackBackground", false);
                run("Make Binary", "thresholded remaining black");
                run("Convert to Mask");

/* Section 7: Identifying peripheral rhino and nuage marker
 */
// creating images of just peripheral rhino and nuage marker.
        imageCalculator("AND create", rhi, nup);
                selectWindow("Result of "+rhi);
                saveAs("Tiff",
output+File.separator+Directory+File.separator+date+"peripheral_raw_"+rhi);
        imageCalculator("AND create", nuage, nup);
                selectWindow("Result of "+nuage);
                saveAs("Tiff",
output+File.separator+Directory+File.separator+date+"peripheral_raw_"+nuage);

// merging thresholded raw data for rhino and nuage marker
        run("Merge Channels...", "c1=["+date+"peripheral_raw_"+nuage+"]
c2="+date+"peripheral_raw_"+rhi+" create keep");
                selectWindow("Composite");
                saveAs("Jpeg",
output+File.separator+Directory+File.separator+date+prefix+"composite_raw.jpg");
                close();

/* Section 8: measuring overlap between Rhino and nuage marker with thresholded data
 */
// preparing rhino and nuage signals for examsion by getting rid of isolated pixels,
// picking and adding Rhino particles to region of interests (ROIs) and
// measruing nuage signal for each RIO with no expansion.
        selectWindow(date+"peripheral_raw_"+rhi);
                run("Close-");
                run("Open");
                run("Analyze Particles...", "size=5-Infinity pixel exclude summarize add");
if (roiManager("count") >= 1){
        run("Set Measurements...", "area mean standard min area_fraction redirect=None
decimal=3");

        selectWindow(date+"peripheral_raw_"+nuage);
                run("Close-");
                run("Open");
                run("Analyze Particles...", "size=5-Infinity pixel exclude summarize");
                roiManager("Measure");
        selectWindow ("Results");
        saveAs("Results",
output+File.separator+Directory+File.separator+date+prefix+"results_ex0.csv");
```

```
        run("Close");

// saving composite image of no expansion
        run("Merge Channels...", "c1=["+date+"peripheral_raw_"+nuage+"]
c2="+date+"peripheral_raw_"+rhi+" create keep");
                selectWindow("Composite");
                saveAs("Jpeg",
output+File.separator+Directory+File.separator+date+prefix+"composite_ex0.jpg");
                close();

/* Section 9: iterations of expanding nuage signal and measuring signal in RIOs
 */
// expansion #1, measure overlap and save composite
        selectWindow(date+"peripheral_raw_"+nuage);
                run("Select None");
                run("Dilate");
                run("Analyze Particles...", "size=5-Infinity pixel exclude summarize");
                roiManager("Measure");
        selectWindow ("Results");
        saveAs("Results",
output+File.separator+Directory+File.separator+date+prefix+"results_ex1.csv");
        run("Close");

        run("Merge Channels...", "c1=["+date+"peripheral_raw_"+nuage+"]
c2="+date+"peripheral_raw_"+rhi+" create keep");
                selectWindow("Composite");
                saveAs("Jpeg",
output+File.separator+Directory+File.separator+date+prefix+"composite_ex1.jpg");
                close();

// expansion #2, measure overlap and save composite
        selectWindow(date+"peripheral_raw_"+nuage);
                run("Select None");
                run("Dilate");
                run("Analyze Particles...", "size=5-Infinity pixel exclude summarize");
                roiManager("Measure");
        selectWindow ("Results");
        saveAs("Results",
output+File.separator+Directory+File.separator+date+prefix+"results_ex2.csv");
        run("Close");

        run("Merge Channels...", "c1=["+date+"peripheral_raw_"+nuage+"]
c2="+date+"peripheral_raw_"+rhi+" create keep");
                selectWindow("Composite");
                saveAs("Jpeg",
output+File.separator+Directory+File.separator+date+prefix+"composite_ex2.jpg");
                close();

// expansion #3, measure overlap and save composite
        selectWindow(date+"peripheral_raw_"+nuage);
                run("Select None");
                run("Dilate");
                run("Analyze Particles...", "size=5-Infinity pixel exclude summarize");
                roiManager("Measure");
        selectWindow ("Results");
        saveAs("Results",
output+File.separator+Directory+File.separator+date+prefix+"results_ex3.csv");
        run("Close");

        run("Merge Channels...", "c1=["+date+"peripheral_raw_"+nuage+"]
c2="+date+"peripheral_raw_"+rhi+" create keep");
                selectWindow("Composite");
                saveAs("Jpeg",
output+File.separator+Directory+File.separator+date+prefix+"composite_ex3.jpg");
                close();

// expansion #4, measure overlap and save composite
        selectWindow(date+"peripheral_raw_"+nuage);
                run("Select None");
                run("Dilate");
                run("Analyze Particles...", "size=5-Infinity pixel exclude summarize");
```

```
                roiManager("Measure");
        selectWindow ("Results");
        saveAs("Results",
output+File.separator+Directory+File.separator+date+prefix+"results_ex4.csv");
        run("Close");

        run("Merge Channels...", "c1=["+date+"peripheral_raw_"+nuage+"]
c2="+date+"peripheral_raw_"+rhi+" create keep");
                selectWindow("Composite");
                saveAs("Jpeg",
output+File.separator+Directory+File.separator+date+prefix+"composite_ex4.jpg");
                close();

// expansion #5, measure overlap and save composite
        selectWindow(date+"peripheral_raw_"+nuage);
                run("Select None");
                run("Dilate");
                run("Analyze Particles...", "size=5-Infinity pixel exclude summarize");
                roiManager("Measure");
        selectWindow ("Results");
        saveAs("Results",
output+File.separator+Directory+File.separator+date+prefix+"results_ex5.csv");
        run("Close");

        run("Merge Channels...", "c1=["+date+"peripheral_raw_"+nuage+"]
c2="+date+"peripheral_raw_"+rhi+" create keep");
                selectWindow("Composite");
                saveAs("Jpeg",
output+File.separator+Directory+File.separator+date+prefix+"composite_ex5.jpg");
                close();

// expansion #6, measure overlap and save composite
        selectWindow(date+"peripheral_raw_"+nuage);
                run("Select None");
                run("Dilate");
                run("Analyze Particles...", "size=5-Infinity pixel exclude summarize");
                roiManager("Measure");
        selectWindow ("Results");
        saveAs("Results",
output+File.separator+Directory+File.separator+date+prefix+"results_ex6.csv");
        run("Close");

        run("Merge Channels...", "c1=["+date+"peripheral_raw_"+nuage+"]
c2="+date+"peripheral_raw_"+rhi+" create keep");
                selectWindow("Composite");
                saveAs("Jpeg",
output+File.separator+Directory+File.separator+date+prefix+"composite_ex6.jpg");
                close();

// save summary file after all espansions for each z section
        selectWindow ("Summary");
        saveAs("Results",
output+File.separator+Directory+File.separator+date+prefix+"summary.csv");
        run("Close");
        }
}
setBatchMode(false)
```
